# SpoIVA contributes to efficient engulfment through a cytoskeletal-like mechanism during *Bacillus subtilis* sporulation

**DOI:** 10.64898/2026.07.08.736982

**Authors:** Betty Fekade, Siham Gabow, Kaitlyn Coleman, Saskia Bakker, Chris L.B. Graham, Cécile Morlot, Christopher D. A. Rodrigues

**Affiliations:** School of Life Sciences, University of Warwick, UK; Univ. Grenoble Alpes, CNRS, CEA, IBS, F-38000 Grenoble, FR

**Keywords:** SpoIVA, Coat, Spores, *Bacillus subtilis*, Engulfment, Phagocytosis

## Abstract

During endospore (spore) development in bacteria, polar cell division generates two transcriptionally distinct cellular compartments, the mother cell and future spore (forespore). Signalling between these cells leads to sequential and compartmentalized transcription, along with key morphogenetics events, including the phagocytic-like process of engulfment and the recruitment of coat proteins to the engulfing membrane. The SpoIVA ATPase is an essential sporulation protein that assembles into static filaments at the forespore surface during engulfment, where it functions as the basement layer for coat assembly. Here, using *Bacillus subtilis,* we reveal an additional role for SpoIVA during engulfment. Cytological analysis of a *spoIVA* null mutant (Δ*spoIVA*) revealed engulfment defects such as septal membrane bulges and asymmetric membrane migration, similar to those typically associated with impaired peptidoglycan remodelling during engulfment. Engulfment defects were exacerbated when Δ*spoIVA* was combined mutants known to impact engulfment progression and efficiency. Importantly, a *spoIVA* mutant (K30A) impaired for ATP hydrolysis and filament formation *in vitro* but partially functional for coat assembly *in vivo*, closely phenocopies the *spoIVA* null mutant engulfment defects. Based on these data, we propose a model whereby SpoIVA polymerisation at the spore surface, independently of coat assembly, plays a mechanical and structural role during engulfment, akin to the cytoskeletal proteins that drive phagocytosis in eukaryotic cells.

**IMPORTANCE:** Endospore formation relies on membrane remodeling to generate highly resistant dormant spores in human, animal and insect pathogens. During engulfment, the mother-cell membrane migrates around the forespore in a phagocytic-like process while the spore coat is simultaneously assembled. SpoIVA is widely recognized as the ATPase that polymerizes at the forespore surface to nucleate assembly of the multilayered spore coat. Here, we demonstrate that SpoIVA also performs a distinct function during engulfment that is independent of its role in coat formation. Cells lacking SpoIVA, or carrying a polymerization-defective SpoIVA variant, exhibit membrane deformations characteristic of impaired engulfment. We propose that SpoIVA polymers provide structural and mechanical support for membrane migration, analogous to the role of cytoskeletal systems during eukaryotic phagocytosis. These findings reveal an unexpected function for SpoIVA and demonstrate that bacterial protein polymers can provide mechanical support for membrane remodeling during complex developmental processes, analogous to cytoskeletal systems in eukaryotic cells

## INTRODUCTION

In response to starvation, or other environmental signals, many bacteria from the *Firmicutes* phylum enter a developmental pathway called sporulation, that results in the formation of a highly resistant, dormant endospore (spore) (1, 2). Sporulation involves various stages and begins with the formation of a polar septum that divides the developing cell in two transcriptionally distinct compartments of different size: a larger cell called the mother cell and smaller one, called the forespore (1, 2). Next, the mother membranes migrate around the forespore in a phagocytic-like process known as engulfment, which results in internalisation of the forespore within the mother cell cytoplasm and the generation of the spore double-membrane envelope: an inner membrane originating from the forespore itself and an outer membrane derived from the mother cell, with a periplasmic-like space in between, containing a thin layer of peptidoglycan called the germ cell wall (3–6). Within the mother cell, the spore develops its protective cellular envelope largely composed of spore-specific peptidoglycan called the cortex and a complex, multilayered coat (4, 7, 8). As the spore reaches maturity, the mother cell lyses, releasing it into the environment where it remains dormant until favourable conditions arise. In this work, we focus on the engulfment process and how SpoIVA, an essential and highly conserved protein known for its role in the formation of the spore envelope (9–11), also contributes to engulfment.

Sporulation is controlled by four, compartment-specific sigma factors, σF, σE, σG, and σK (1, 2). σF and σE are active during the early stages of spore development, in the forespore and mother cell, respectively, while σG and σK become active after engulfment completion in the forespore and mother, respectively (1, 2). σE plays an important role in development, as it drives the production of proteins that orchestrate the engulfment process, as well as many of the proteins that build the spore coat, including SpoIVA (12).

Current data indicate that the phagocytic-like process of engulfment is controlled by multiple processes, including peptidoglycan (abbreviated as PG) hydrolysis and synthesis, membrane synthesis, and a conserved intercellular protein interaction between SpoIIQ in the forespore and SpoIIIAH in the mother cell (known as the SpoIIIAH-SpoIIQ ratchet). PG hydrolysis is primarily controlled by the DMP complex, named after its constituents, SpoIID, SpoIIM, and SpoIIP, which localize at the polar septum and are required for septal PG thinning (5, 13, 14). PG synthesis is also required for engulfment; however, the exact PG synthetic enzymes remain unidentified (3, 15, 16). The SpoIIIAH-SpoIIQ ratchet, by bridging the mother cell and forespore membranes, contributes to migration of the mother cell membrane around the forespore (17). While DMP complex activity is absolutely required for initiation of engulfment, PG synthesis and the SpoIIIAH-SpoIIQ ratchet contribute to its efficiency, allowing for symmetric and timely envelopment of the forespore by the mother cell (16).

Importantly, as engulfment progresses, key morphogenetic proteins required for coat assembly become localized in the engulfing membrane, initiating the ordered assembly of the coat from its innermost to outermost layer (8, 11). The basement layer, the coat’s innermost region, anchors subsequent layers to the spore outer membrane and is formed by SpoVM and SpoIVA (8, 11). SpoVM is a small protein composed of a single, amphipathic, α-helix peptide anchored into the outer forespore membrane through hydrophobic interactions (18). SpoVM recognizes the positive outer membrane curvature on the mother cell side during the early stages of engulfment and interacts with SpoIVA at this location (18, 19). SpoIVA is an ATPase that is universally conserved in spore-formers and polymerises into filament-like structures around the spore (10, 20). In the absence of SpoIVA, attachment of the coat layers fails, resulting in their accumulation in the mother cell cytoplasm (9). Furthermore, the cortex PG fails to assemble due to mislocalisation of SpoVK, a chaperone of MurG, involved in synthesis of the PG precursor, Lipid II (21). Similarly, in the absence of SpoVM, the spore cortex is absent and the coat is partially attached (22). Assembly of the inner coat requires SpoVID and SafA, which depend on SpoIVA for their localisation to the forespore (11, 23, 24). The outer coat is assembled by CotE, another morphogenetic protein whose localization requires SpoIVA and SpoVID (25). Initially, these coat morphogenetic proteins localize as an organised scaffold on the mother cell-proximal pole of the forespore. As engulfment completes, they encircle the spore in a process called encasement, driven by SpoVM and SpoVID (26, 27).

Interestingly, while mutants affecting engulfment have been reported to impact coat localisation and assembly (11, 28), it remains unclear whether coat assembly or coat proteins contribute to engulfment. Here, building on our previous work on the role of spore envelope layers in spore shape (29), we uncovered a previously unappreciated role for SpoIVA during engulfment. Our data indicate that sporangia lacking SpoIVA exhibit engulfment defects similar to those exhibited by hypomorphic alleles of DMP complex genes. Moreover, sporangia lacking SpoIVA and MurAB, which is required for efficient PG precursor synthesis during engulfment (16), exhibit more pronounced engulfment defects compared to either single mutant. Importantly, a *spoIVA* mutant (K30A) that is unable to hydrolyse ATP and fails to assemble into static filaments at a critical concentration *in vitro* (20), yet partially supports coat assembly *in vivo* (20), closely phenocopies the engulfment defects of the *spoIVA* null mutant. Collectively, these data support a role for SpoIVA during engulfment. We propose a model whereby SpoIVA polymerisation, independently of its function in coat assembly, contributes to membrane scaffolding during engulfment, akin to the role of cytoskeletal proteins during the formation of the phagocytic cup in eukaryotic cells.

## RESULTS

### SpoIVA is required for efficient initiation and completion of engulfment

During our analysis of how the spore envelope contributes to spore shape (29), we noticed that the Δ*spoIVA* mutant resulted in engulfment defects in a subset of cells. To more precisely determine the extent of these defects, we examined wild-type (WT) and Δ*spoIVA* strains at 2 hours onset of sporulation (T2). Consistent with previous results, most WT cells harboured a curving septal membrane that evenly began to surround the forespore. In contrast, in the Δ*spoIVA* mutant, the septal membrane often contained a small “bulge” protruding into the mother cell (Fig. 1A-C). Furthermore, some Δ*spoIVA* sporangia exhibited uneven septal membrane migration around the forespore (Fig. 1B). Quantification at T2 confirmed the higher proportion of septal bulges (38.2%) and uneven membrane migration (16.9%) in the Δ*spoIVA* mutant compared to WT cells (Fig. 1D). We also examined WT and Δ*spoIVA* cells at T3, when most cells are approaching engulfment completion (16). In most WT and Δ*spoIVA* cells, membrane migration appeared complete or near complete (Fig. 1B). However, among cells still undergoing engulfment, the Δ*spoIVA* mutant exhibited a higher proportion of septal bulges and uneven membrane migration compared to WT (Fig. 1D). At T3, we observed an additional phenotype in Δ*spoIVA* cells that had completed engulfment: “indented” forespores (9.1% of all Δ*spoIVA* sporangia), characterized by an indentation along the long cell axis, resulting in a deformed forespore shape (Fig. 1B & E).

**Figure 1.**
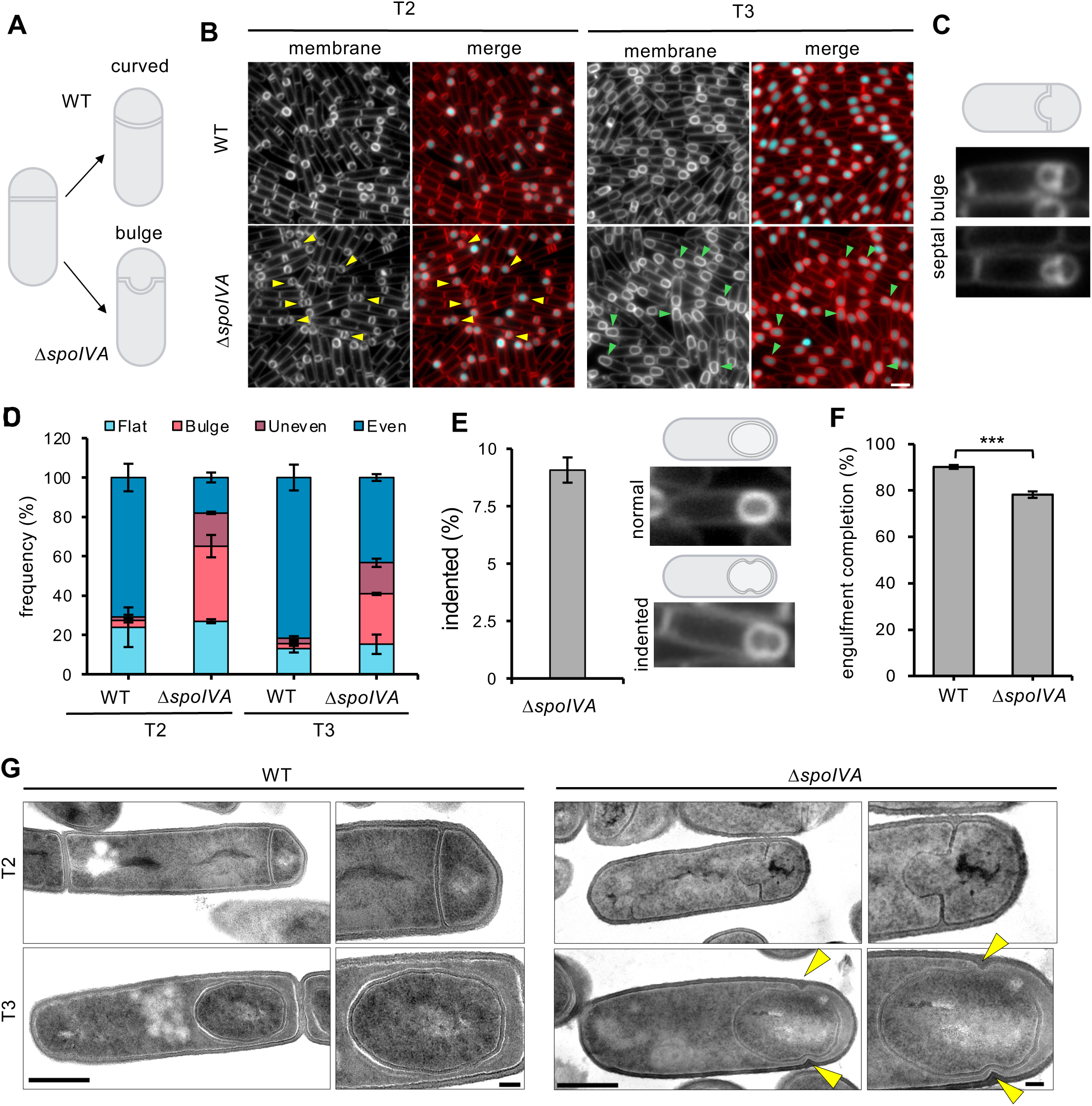
Engulfment defects in the Δ*spoIVA* mutant. **(A)** Schematic representation of normal engulfment in a wild-type (WT) cell (top) and abnormal engulfment in a Δ*spoIVA* mutant cell (bottom). **(B)** Engulfment progression in the wild-type (WT) and Δ*spoIVA* strains at 2 hours (T2) and 3 hours (T3) after the onset of sporulation. Forespore cytoplasm were visualised using a forespore reporter (P*spoIIQ*-cfp) that appears cyan in merged images. Cell membranes are false-coloured red in merged images. Septal membrane bulges are highlighted with yellow triangles and indented cells are highlighted with green triangles. Scale bar = 2 μm. **(C)** Representative image of bulge septa in Δ*spoIVA* mutants at T2, more representative images are depicted in Fig.S6. **(D)** Average frequency (mean percentage ± SD, n=3) of sporulating cells with flat (light blue), bulging (pink), uneven (purple) and even (dark blue) septa during a sporulation time course in WT and Δ*spoIVA* cells (n>100 per time point, per strain and per replicate). Error bars indicate the standard deviation from three biological replicates. Representative images of cells with each of the septal phenotypes are depicted in Fig. S10. **(E)** Mean frequency (mean percentage ± SD n=3) of indented cells relative to normal cells in WT and Δ*spoIVA* cells at T3. Error bars indicate the SD from three biological replicates. Schematic representation of normal and indented cells at T3, with representative images of phenotype. \*\*\**p* <0.001 using Welch’s t test performed on the mean of replicates (n=3) at T3 for the WT versus the Δ*spoIVA* mutant. **(F)** Mean frequency (mean percentage ± standard deviation [SD], n=3) of engulfment completion in WT and Δ*spoIVA* cells at T3. \*\*\**p* <0.001 using Welch’s t test performed on the mean of replicates (n=3) at T3 for the WT versus the Δ*spoIVA* mutant. **(G)** Representaitve images of sporulating cells in the WT, Δ*spoIVA* mutant at 2 hours (T2) and 3 hours (T3) after the onset of sporulation. Δ*spoIVA* mutant bulging phenotype at T2 and indented phenotype after engulfment completion at T3. Scale bar is 500 nm and 100 nm in standard and zoomed-in images, respectively.

To assess whether these engulfment defects affected the efficiency of engulfment completion, we analysed forespores at T4, when 80-90% of WT sporangia have typically completed engulfment (16). In the Δ*spoIVA* mutant, only 78.3% of sporangia had completed engulfment by T4, compared to 90.2% in the WT (Fig. 1F). To rule out specific effects due to the antibiotic resistance marker, we compared spore morphogenesis of Δ*spoIVA* strains containing different antibiotic markers and confirmed that these observed phenotypes were due to the absence of SpoIVA (Fig. S1C). Furthermore, complementation of SpoIVA at an ectopic locus reversed the morphological defects of the Δ*spoIVA* mutation (Fig. S1A & B).

Finally, as SpoIVA localization partially depends on SpoVM (Fig. S2A) (18), we tested if the Δ*spoVM* mutant exhibited similar engulfment defects. While septal bulges and indented forespores were observed in Δ*spoVM*, these phenotypes appeared less frequent than in Δ*spoIVA* (Fig. S2B). We conclude that the partial mislocalization of SpoIVA in the absence of SpoVM results in a less penetrant engulfment phenotype than that observed in the absence of SpoIVA.

Taken together, these results indicate that SpoIVA is required for efficient initiation and completion of engulfment.

### Indented forespores in the Δ*spoIVA* mutant result from septal peptidoglycan protrusions

To examine the morphological defects of the *ΔspoIVA* mutant in greater detail, we used transmission electron microscopy (TEM) to analyse sporulating cells at T2 and T3. At T2, we identified forespores with septal bulges protruding into the mother cell (Fig.1G and S3). These bulges were delineated by septal material perpendicular to the long axis of the sporangia, likely resulting from incomplete thinning of the septum. These phenotypes are reminiscent of those observed in DMP complex and SpoIIB mutants (5, 30–32). At T3, we also identified sporulating cells with indented forespores (Fig.1G and S3), where the indentations resulted from mother cell PG material protruding towards the forespore. Given the similarity in the position of these protrusions to the polar septum, it is likely that they represent incompletely digested septa.

Together, these data suggest that the absence of SpoIVA compromises the efficiency of septal PG thinning, leading to incomplete hydrolysis that results in septal bulges early in engulfment, and septal protrusions that deform forespore morphology once engulfment is complete.

### SpoIIP localization is heterogeneous in the absence of SpoIVA

Our data suggest that SpoIVA is required for efficient engulfment. One way in which SpoIVA could affect engulfment efficiency is by affecting the localization of the DMP complex. To test this, we examined the localization of GFP-SpoIIP, which localizes to the septal membrane and remains localized at the leading edge of the engulfing membrane throughout engulfment (33).

Consistent with previous reports (16, 33), in WT cells, GFP-SpoIIP was faintly present in the mother cell membranes and was enriched in septal membranes initiating engulfment or formed bright foci at the leading edge of the engulfing membrane in sporangia progressing through engulfment (Fig. 2A). In the Δ*spoIVA* mutant, GFP-SpoIIP exhibited similar localization patterns, although with more diffuse signal in the mother cell (Fig. 2A).

**Figure 2.**
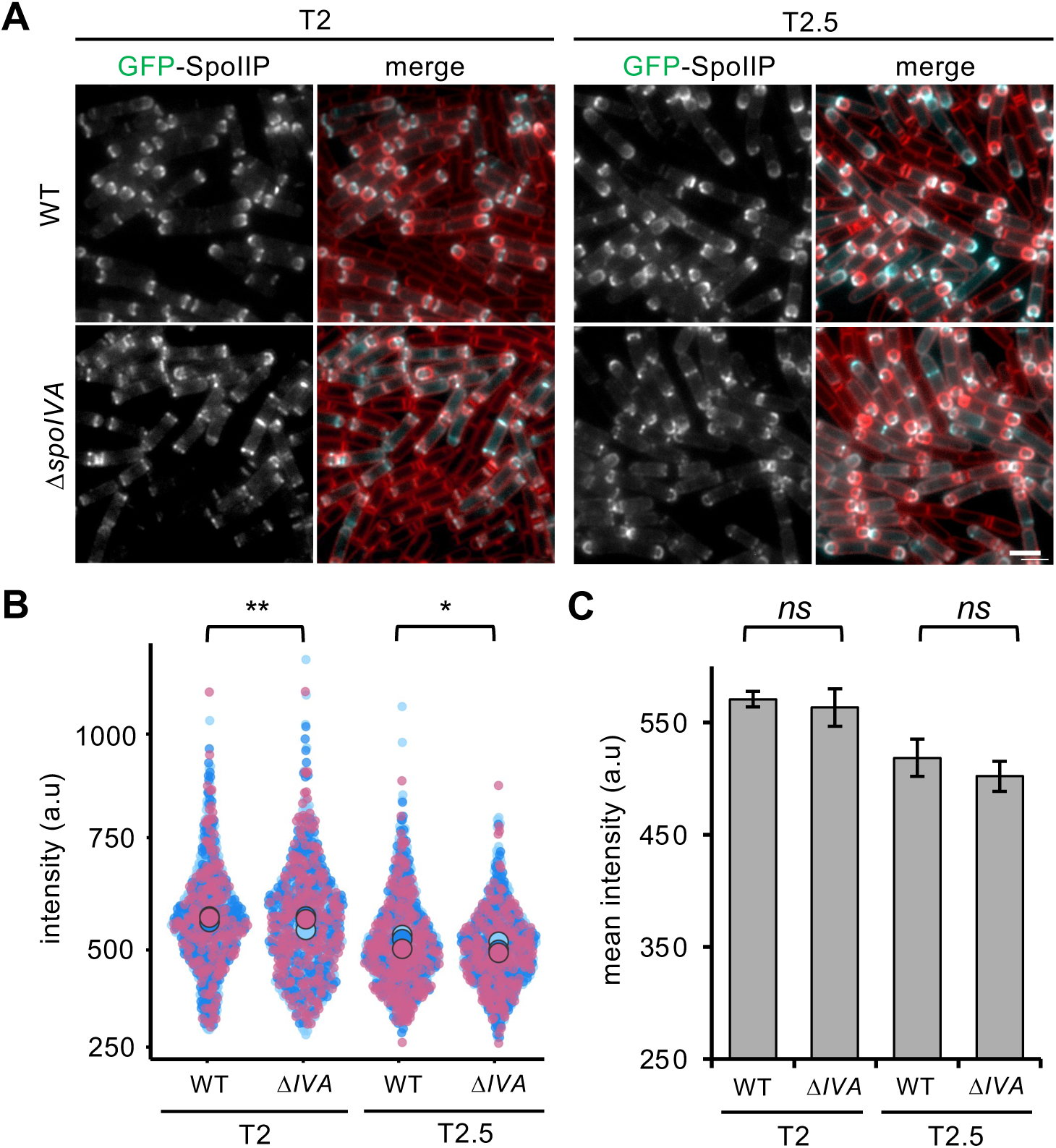
GFP-SpoIIP localization in the absence of SpoIVA. **(A)** GFP-SpoIIP localisation in wild-type (WT) and Δ*spoIVA* strain. Cell membranes and GFP are false-coloured red and cyan, respectively, in merged images. Scale bar = 2 μm **(B)** Distribution of GFP-SpoIIP intensity. *\*p* < 0.05 and \*\*\**p* < 0.01 using Kolmogorov-Smirnov test to compare total population distributions. Data from 3 biological replicates and a total of 828 and 805 cells for WT and Δ*spoIVA,* respectively, at T2 and 746 and 794 cells for WT and Δ*spoIVA,* respectively, at T2.5. **(C)** Mean GFP-SpoIIP intensity (a.u.) in the WT and Δ*spoIVA* strains 2 hours and 2.5 hours after onset of sporulation. *ns* (not significant) using Welch’s t test performed on the mean GFP-SpoIIP intensity, replicates (n=3) at T2 and T2.5 for the WT versus Δ*spoIVA*.

Quantification of GFP-SpoIIP signal in the septal membrane showed similar average fluorescence intensities in WT and Δ*spoIVA* (Fig. 2B & C). However, analysis of individual cells revealed a broader distribution of GFP-SpoIIP signal intensity in the Δ*spoIVA* mutant at T2 (Fig. 2B). Thus, while SpoIVA is not a major determinant of GFP-SpoIIP localisation, its absence results in more heterogeneous GFP-SpoIIP localization.

### The absence of SpoIVA partially suppresses defects caused by imbalances in PG synthesis and hydrolysis

To further test a role of SpoIVA in promoting PG hydrolysis, we took advantage of a recently described double mutant, Δ*spoIIIM* Δ*pbpG,* which exhibits imbalanced PG remodelling during early engulfment (34). SpoIIIM (a mother cell LysM-domain containing protein) and PbpG (a forespore Class A Penicillin-Binding-Protein) are thought to counteract the activity of the DMP complex by contributing to protein-PG interactions and PG synthesis, respectively (34). In their absence, the septal chromosome translocation pore enlarges, resulting in forespore cytoplasm leakage and miscompartmentalisation during the early stages of engulfment, in approximately 90% of sporangia (34). Importantly, compartmentalisation can be restored in the Δ*spoIIIM* Δ*pbpG* mutant if the efficiency of PG hydrolysis is reduced, such as in the Δ*spoIIB* mutant (34).

If SpoIVA promotes PG hydrolysis, its absence might improve compartmentalization in the Δ*spoIIIM* Δ*pbpG* background. To test this, we compared compartmentalisation in Δ*spoIIIM* Δ*pbpG* and Δ*spoIIIM* Δ*pbpG* Δ*spoIVA* strains at T3 using a forespore-produced fluorescent transcriptional reporter (P*_spoIIQ_*-*cfp*). Consistent with previous results (34), in the Δ*spoIIIM* Δ*pbpG* double mutant, 89.3% of sporangia harboured miscompartmentalised forespores (Fig. 3A, 3C & S4). In the Δ*spoIIIM* Δ*pbpG* Δ*spoIVA* triple mutant, miscompartmentalisation decreased significantly to 77.8%.

**Figure 3.**
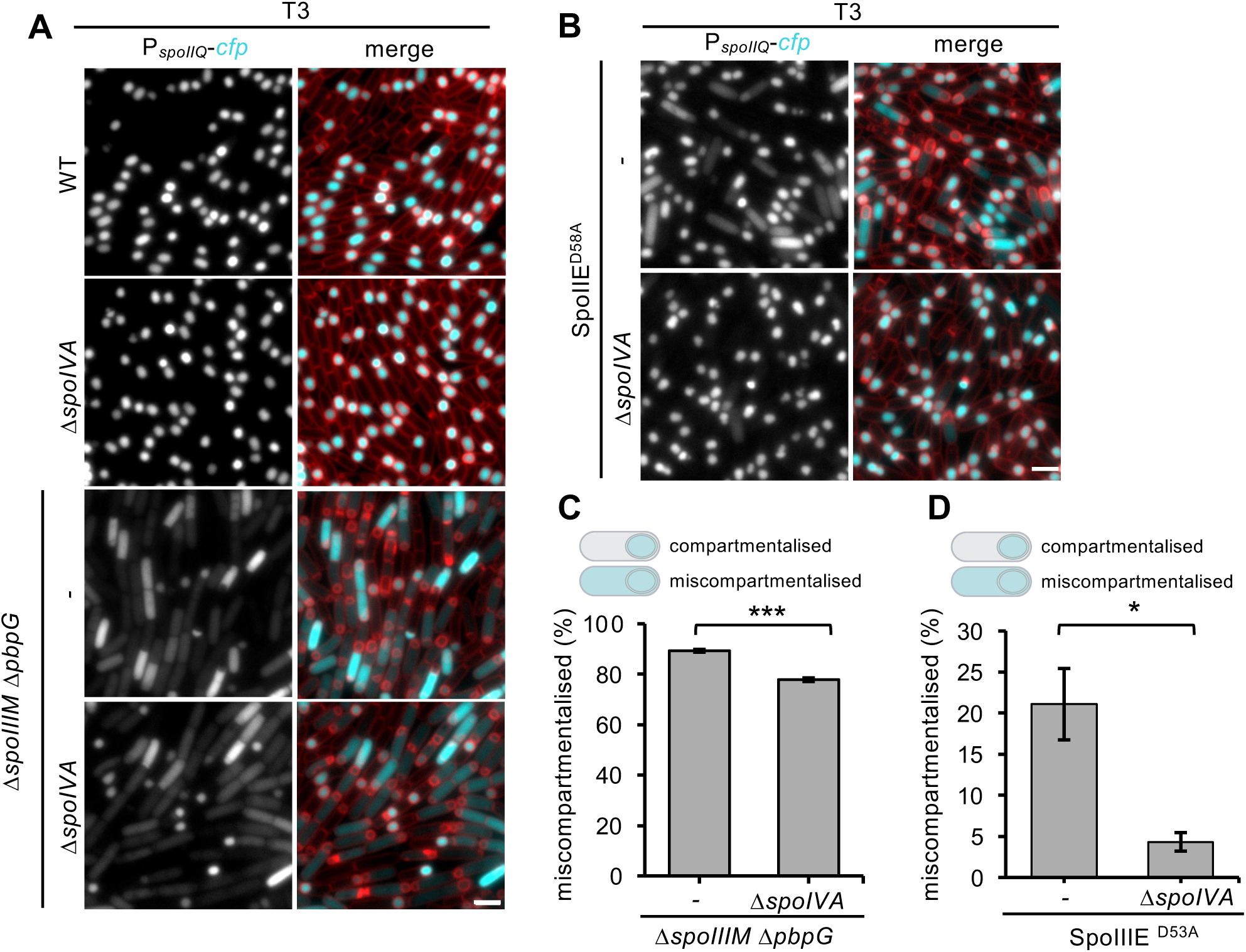
The absence of SpoIVA increases compartmentalisation defects. **(A)** Representation images of misscompartmentalisation in wild-type (WT), Δ*spoIIIM* Δ*pbpG* and Δ*spoIIIM* Δ*pbpG* Δ*spoIVA* strains at 3 hours (T3) after the onset of sporulation. **(B)** Representation images of misscompartmentalisation in SpoIIIE^D58A^, and SpoIIIE^D58A^ Δ*spoIVA* strain at 3 hours (T3) after the onset of sporulation. Forespore cytoplasm were visualised using a forespore reporter (P*_spoIIQ_*-cfp) that appears cyan in merged images. Cell membranes false-coloured red in merged images. Scale bar = 2 μm. **(C)** Average frequency (mean percentage ± SD, n=3) of miscompartmentalisation defect WT, Δ*spoIIIM* Δ*pbpG* and Δ*spoIIIM* Δ*pbpG* Δ*spoIVA* strains cells (n>100 per time point, per strain and per replicate). Error bars indicate the standard deviation from three biological replicates. Schematic representation of normal and miscompartmentalised cells at T3. \*\*\**p* <0.001 by Welch’s t test performed on the mean of replicates (n=3) at T3 for the WT versus the Δ*spoIVA* Δ*murAB* mutant. **(D)** Average frequency (mean percentage ± SD, n=3) of miscompartmentalisation defect SpoIIIED58A, and SpoIIIED58A Δ*spoIVA* strains cells (n>100 per time point, per strain and per replicate). Error bars indicate the standard deviation from three biological replicates. Schematic representation of normal and miscompartmentalised cells at T3. \**p* <0.05 by Welch’s t test performed on the mean of replicates (n=3) at T3 for the WT versus the Δ*spoIVA* Δ*murAB* mutant.

We also tested whether the absence of SpoIVA suppresses miscompartmentalisation in the SpoIIIE^D584A^ mutant, which exhibits miscompartmentalisation in approximately 20% of sporangia (34). At T3, deletion of *spoIVA* in the SpoIIIE^D584A^ background reduced miscompartmentalisation frequency fivefold (from 21.1% to 4.3%) (Figs. 3B and 3D).

Taken together, these data suggest that the absence of SpoIVA reduces the efficiency of PG hydrolysis, thereby suppressing miscompartmentalisation defects associated with imbalances in PG synthesis and hydrolysis.

### The absence of SpoIVA exacerbates the engulfment defects in the *murAB* mutant

To further validate the role of SpoIVA in engulfment, we tested whether deletion of *murAB*, a *murAA* homologue involved in the first step of PG precursor synthesis, would exacerbate the engulfment defects of the Δ*spoIVA* mutant. The absence of MurAB leads to reduced PG precursor synthesis, leading to slower engulfment and mild mother cell membrane migration defects (16).

We compared the Δ*spoIVA* and Δ*spoIVA* Δ*murAB* mutants at T2, when Δ*spoIVA* engulfment defects are more pronounced (Fig. 4A). Quantification showed that the Δ*spoIVA* Δ*murAB* double mutant produced more septal bulges (57.1%) than the Δ*spoIVA* single mutant (44.9%) (Fig. 4B). At T3, the Δ*spoIVA* Δ*murAB* double mutant also showed a significant decrease in engulfment completion efficiency compared to the single mutants (Fig. 4D & S5). Consistent with the increase in septal bulges, we observed a higher frequency of indented forespore at T3 in the Δ*spoIVA* Δ*murAB* double mutant (18.6%) (Fig. 4C & S5), relative to Δ*spoIVA* (9.1%) (Fig.1E)

**Figure 4.**
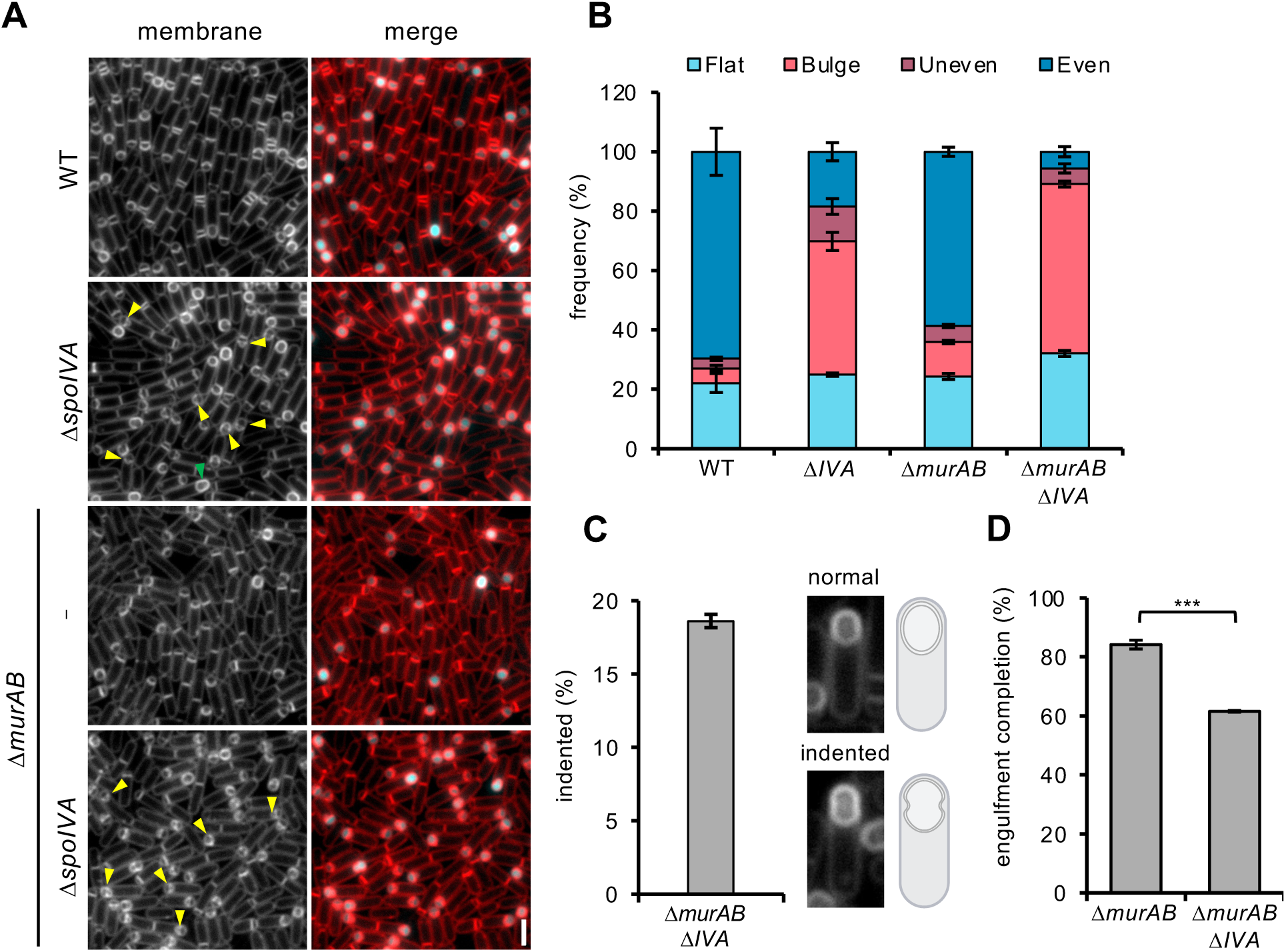
Engulfment defects in the Δ*spoIVA, ΔmurAB and Δ spoIVA ΔmurAB* mutant. **(A)** Engulfment progression in the wild-type (WT), Δ*spoIVA,* Δ*murAB and* Δ*spoIVA* Δ*murAB* strains at 2 hours (T2) after the onset of sporulation. Forespore cytoplasm were visualised using a forespore reporter (P*_spoIIQ_*-cfp) that appears cyan in merged images. Cell membranes are false-coloured red in merged images. Septal membrane bulges are highlighted with yellow triangles, and indented cells are highlighted with green triangles. Scale bar = 2 μm. **(B)** Average frequency (mean percentage ± SD, n=3) of sporulating cells with flat (light blue), bulging (pink), uneven (purple) and even (dark blue) septa during a sporulation time course in WT Δ*spoIVA, ΔmurAB and Δ spoIVA ΔmurAB* mutant cells (n>100 per time point, per strain and per replicate). Error bars indicate the standard deviation from three biological replicates. Representative images of cells with each of phenotype are depicted in Fig. S10. **(C)** Mean frequency (mean percentage ± SD, n=3) of indented cells relative to normal cells, in WT and Δ*spoIVA* at T3. Error bars indicate the SD from three biological replicates. Schematic representation of normal and indented cells at T3, with representative images of phenotype. \*\*\**p* <0.001 using Welch’s t test performed on the mean of replicates (n=3) at T3 for the WT versus the Δ*spoIVA* Δ*murAB* mutant. **(D)** Mean frequency (mean percentage ± SD, n=3) of engulfment completion in Δ*spoIVA* and *ΔspoIVA ΔmurAB* cells at T3. \*\*\**p* <0.001 using Welch’s t test performed on the mean of replicates (n=3) at T3 for the WT versus the Δ*murAB* mutant and WT vs Δ*spoIVA* Δ*murAB* mutant.

Since the absence of MurAB results in a subtle but reproducible decrease in GFP-SpoIIP localisation (16), we examined whether the Δ*spoIVA* Δ*murAB* double mutant caused more severe GFP-SpoIIP localisation defects (Fig. S6A). Quantification confirmed that GFP-SpoIIP signal in the engulfing membrane was lower in the Δ*spoIVA* Δ*murAB* double mutant than in the single mutants (Fig. S6B-E).

These data suggest the absence of SpoIVA exacerbates engulfment defects associated with reduced PG synthesis.

### SpoVK is not required for efficient engulfment

As the above data suggest that the absence of SpoIVA exacerbates the engulfment defects of the *murAB* mutant, we considered the possibility that SpoIVA may influence PG synthesis during engulfment. One way it could do so is through its role in the localisation of SpoVK, a recently identified chaperone that promotes Lipid II synthesis by MurG and depends on SpoIVA for its localisation near the spore membrane (21). To test whether engulfment defects in Δ*spoIVA* were due to SpoVK mislocalisation, we compared the WT, Δ*spoVK* and Δ*spoIVA* strains at T2 and T3.5 (Fig. S7). The Δ*spoVK* mutant showed engulfment membrane migration patterns similar to WT cells and lacked the phenotypes described thus far for Δ*spoIVA.* We conclude that SpoVK is not required for efficient engulfment and that the engulfment defects associated with the absence of SpoIVA are not an indirect consequence of SpoVK mislocalisation.

### Sporulating cells lacking SpoIVA and SpoIIQ exhibit a severe engulfment defect

Efficient engulfment requires the SpoIIIAH-SpoIIQ ratchet, which promotes forward migration of the mother cell membrane around the forespore (17). The role of this ratchet becomes particularly critical when other aspects of engulfment, such as PG synthesis, are compromised (16). To test whether SpoIVA and SpoIIIAH-SpoIIQ have additive roles in engulfment, we examined the Δ*spoIIQ* Δ*spoIVA* double mutant at T2, T3 and T4, comparing it to the Δ*spoIIQ* single mutant.

Consistent with previous reports (35–37), Δ*spoIIQ* forespores were smaller than WT, and by T4, the majority of Δ*spoIIQ* sporangia had completed engulfment (Fig. S8). In the Δ*spoIIQ* Δ*spoIVA* mutant however, engulfment was severely compromised: by T4, the majority of sporangia had failed to complete membrane migration around the forespore. Furthermore, at T4, while some sporangia contained septa that remained at the initial curving stage of engulfment, a high proportion still contained flat polar septa (Fig. S8). Thus, the combined loss of SpoIVA and the SpoIIIAH-SpoIIQ ratchet results in a severe engulfment defect, further supporting the idea that SpoIVA plays a role in engulfment.

### Polymerization of SpoIVA is required for its role in engulfment

SpoIVA is essential for spore coat assembly; in its absence, the coat accumulates in the mother cell cytoplasm rather than encasing the forespore (9). To test whether the engulfment defects in the Δ*spoIVA* mutant are an indirect consequence of coat misassembly, we analyzed the *spoIVA^K30A^* mutant, which can localize the encasement protein SpoVID and outer coat protein CotE (Fig. S9) but is defective in ATP hydrolysis and filament polymerization *in vitro* (20). SpoIVA K30A harbours a disrupted Walker A motif and leads to a 20-fold decrease in sporulation efficiency, compared to the complete sporulation block in Δ*spoIVA* cells (20) .

We compared the Δ*spoIVA* and *spoIVA K30A* mutants at T2 and T3 and found that they closely phenocopied each other (Fig. 5A). At T2, the *spoIVA K30A* mutant displayed septal bulges, but in lower proportion compared to the Δ*spoIVA* mutant, 26.8% and 39.6%, respectively (Fig. 5B). Furthermore, the *spoIVA K30A* mutant also exhibited uneven membrane migration but again at a lower frequency than the Δ*spoIVA* mutant, 17.3% and 11.4%, respectively (Fig. 5B). Furthermore, consistent with the results shown earlier for Δ*spoIVA*, the proportion of septal bulges was reduced at T3 in both mutants (10% in Δ*spoIVA*, 14% in *spoIVA K30A*) (Fig. 5B). Indented forespores were also present in the *spoIVA K30A* mutant, though at a lower frequency (3.3%) than in Δ*spoIVA* (9.1%) (Fig. 5C). Finally, by T4, fewer sporangia had completed engulfment in the *spoIVA K30A* mutant (82.8%) compared to WT (90.2%) (Fig. 5D).

**Figure 5.**
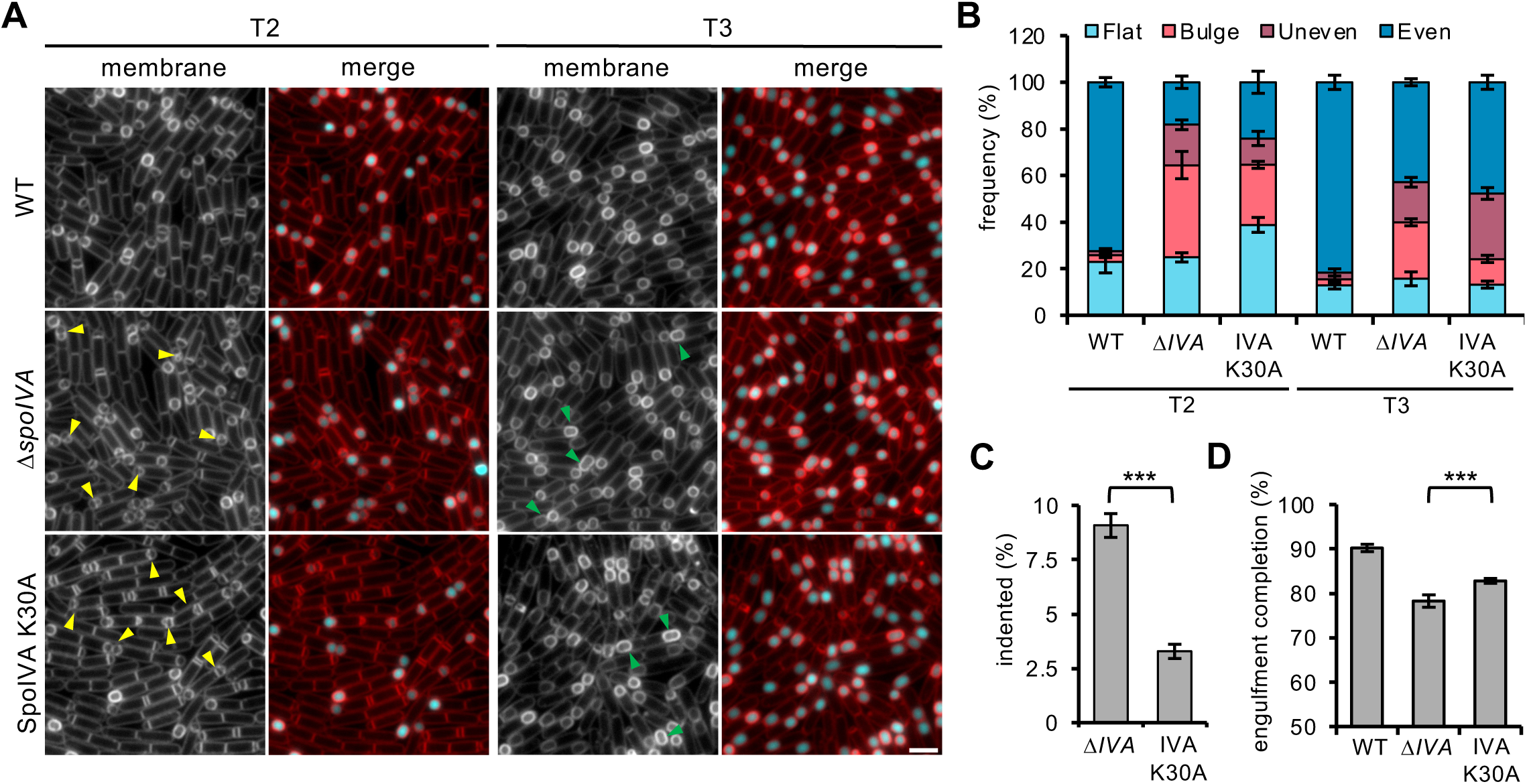
Engulfment defects in the Δ*spoIVA and* SpoIVA-K30A mutant. **(A)** Engulfment progression in the wild-type (WT), Δ*spoIVA,* SpoIVA-K30A strains at 2 hours (T2) after the onset of sporulation. Forespore cytoplasm were visualised using a forespore reporter (P*_spoIIQ_*-cfp) that appears cyan in merged images. Cell membranes are false-coloured red in merged images. Septal membrane bulges are highlighted with yellow triangles and indented cells are highlighted with green triangles. Scale bar = 2 μm. **(B)** Mean frequency (mean percentage ± SD, n=3) of sporulating cells with flat (light blue), bulging (pink), uneven (purple) and even (dark blue) septa during a sporulation time course in WT, Δ*spoIVA* and SpoIVA K30A mutant cells (n>100 per time point, per strain and per replicate). Error bars indicate the standard deviation from three biological replicates. Representative images of cells with each of the septal phenotypes are depicted in Fig. S10. **(C)** Mean frequency (mean percentage ± SD, n=3) of indented cells relative to normal cells, in WT and Δ*spoIVA* cells at T3. Error bars indicate the SD from three biological replicates. \*\*\**p* <0.001 using Welch’s t test performed on the mean of replicates (n=3) at T3 for the WT versus the SpoIVA K30A mutant. **(D)** Mean frequency (mean percentage ± SD, n=3) of engulfment completion in Δ*spoIVA* and SpoIVA K30A cells at T3. \*\*\**p* <0.001 using Welch’s t test performed on the mean of replicates (n=3) at T3 for the WT versus the Δ*spoIVA* mutant and WT vs SpoIVA K30A mutant.

Collectively, these results suggest that the engulfment defects in the Δ*spoIVA* mutant are not a consequence of coat misassembly but instead reflect a direct role for SpoIVA in engulfment. Furthermore, since the SpoIVA K30A mutation impacts ATP hydrolysis and filament formation, these data suggest that ATP hydrolysis and/or SpoIVA polymerization are required for its function in engulfment.

## DISCUSSION

SpoIVA is one of the most well-characterized sporulation proteins, but its role in engulfment has been missed. Here, using cytological analyses, we reveal that SpoIVA plays a direct role during engulfment. Our data show that the absence of SpoIVA results in mild engulfment defects, such as septal bulging, and that these defects are further exacerbated in mutants displaying moderate engulfment impairments. Furthermore, our findings suggest that SpoIVA influences the efficiency of PG hydrolysis without substantially affecting the localisation of the DMP complex. Although GFP-SpoIIP retained most of its localization in the engulfing membrane in the absence of SpoIVA (Fig. 2), septal PG hydrolysis efficiency decreased, which led to partial suppression of miscompartmentalisation defects associated with unbalanced PG remodelling (Fig. 3) and increased engulfment defects (Fig. 4). Below, we discuss possible roles for SpoIVA during engulfment.

While our data indicate that the absence of SpoIVA has a subtle effect on SpoIIP localisation (Fig. 2), we propose that these differences alone are unlikely to fully account for the engulfment defects observed in the Δ*spoIVA* mutant. Supporting this idea, the SpoIVA K30A mutant, which localizes to the spore surface but is affected in its ATP hydrolysis and polymerization abilities (20), still exhibited engulfment defects similar to those observed in the absence of SpoIVA (Fig. 5). Thus, SpoIVA ATP hydrolysis and/or filament formation at the forespore surface may contribute to efficient septal PG thinning and membrane migration around the forespore during engulfment. This interpretation is further supported by recent data by Bauda *et al*., (45) showing that SpoIVA dimerization, which is required for proper SpoIVA localization and, consequently, filament positioning, is critical for SpoIVA function, as dimerization-defective mutants also exhibit engulfment defects similar to the ones we describe here. Altogether, these observations support the idea that SpoIVA polymerisation on the forespore surface facilitates efficient engulfment.

How might SpoIVA polymerisation influence engulfment? Recent *in-situ* cryo-electron tomography data by Bauda *et al,* (45) revealed that SpoIVA assembles into highly organized polymers at the surface of the outer forespore membrane, forming tracks that run parallel to the direction of the engulfing membranes. Although the functional significance of these structures remains unclear, we propose two potential roles: 1) SpoIVA polymers may provide mechanical reinforcement to maintain membrane shape at the onset of septal PG thinning, and 2) the tracks formed by these polymers may structurally confine the localisation of membrane proteins within the outer forespore membrane (Fig. 6). These proposed roles draw parallels with the process of phagocytosis in eukaryotic cells.

**Figure 6.**
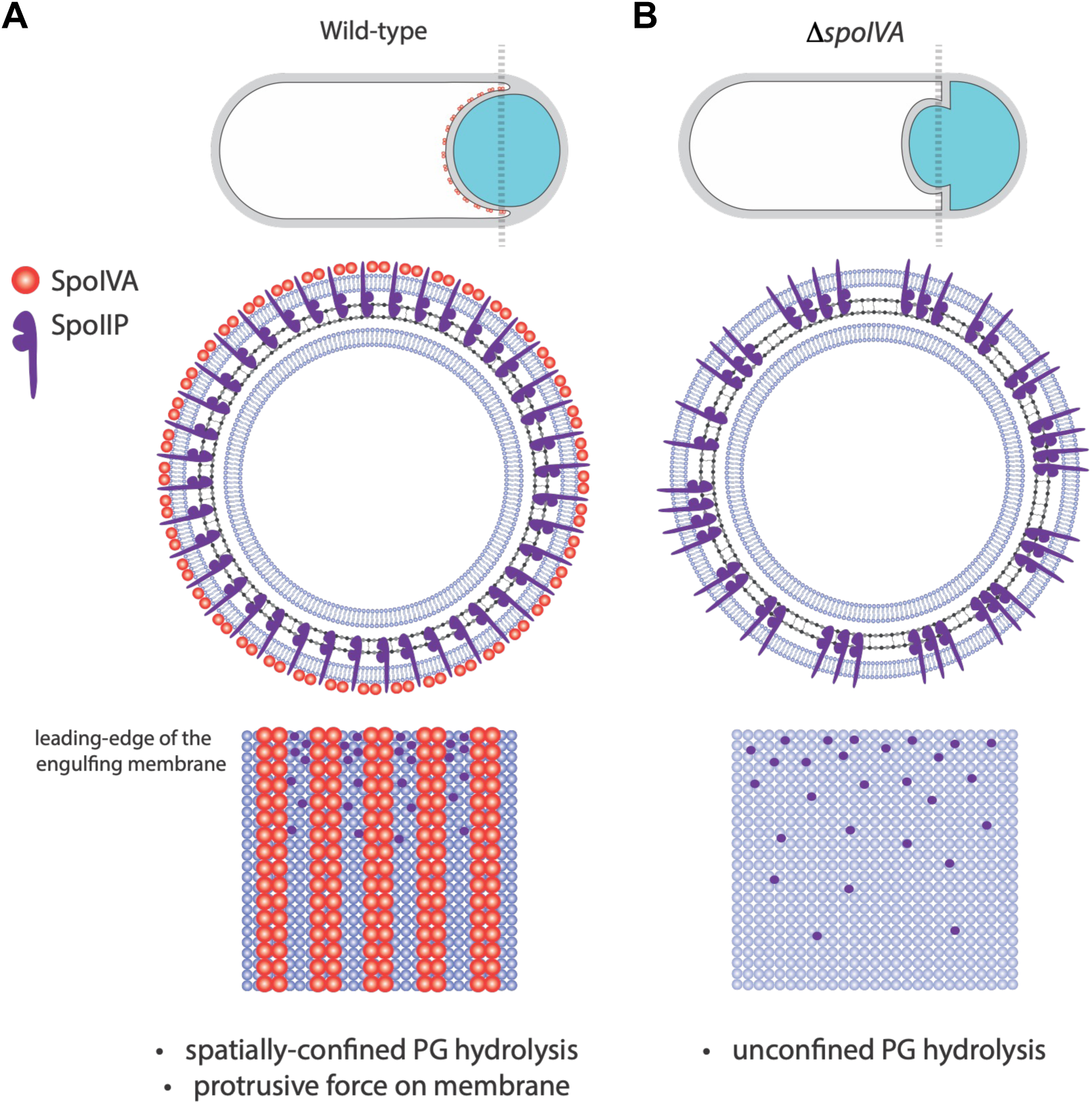
Model illustrating a hypothetical role for SpoIVA during engulfment. **(A)** In WT cells, SpoIVA filaments form tracks that provide mechanical support to the engulfing membrane, pushing it forward. These SpoIVA tracks also confine the localization of the DMP complex (for simplicity only SpoIIP is shown), promoting localized and efficient PG hydrolysis. This two-in-one function for SpoIVA promotes uniform migration of the engulfing membrane and efficient engulfment. **(B)** In cells lacking SpoIVA, the engulfing membrane exhibits reduced mechanical support and less efficient forward membrane migration. In the absence of SpoIVA tracks, DMP complex localization is less confined, leading to less efficient PG hydrolysis, incomplete septal PG thinning, septal bulges and septal remnants. Top panel illustrates the shape of engulfing membrane. The middle panel illustrates the leading edge of the engulfing membrane as seen from a transverse plane. The bottom illustrates a section of the mother cell engulfing membrane, with SpoIVA tracks and the N-terminal domain of SpoIIP seen on the membrane surface.

During phagocytosis, the formation of the phagocytic cup relies on dynamic cytoskeletal proteins, including F-actin and various myosin motors, along with regulatory factors (38). F-actin plays both mechanical and structural roles during phagocytosis: its polymerization generates protrusive forces that drive plasma membrane expansion (39), and in conjunction with proteins like CD44, it confines the movement of membrane-bound proteins (e.g. phagocytic receptors) to create specialized domains of activity (40). Although F-actin undergoes treadmilling and SpoIVA does not, the tracks formed by SpoIVA at the forespore surface may serve analogous function during engulfment. Specifically, SpoIVA tracks could dictate the positioning of the DMP machinery at the leading edge of the engulfing membrane, ensuring coordinated and directional PG hydrolysis and membrane migration. Given that SpoIID degrades PG glycan strands stripped of their stem peptides by SpoIIP, and that SpoIIP activity is enhanced by SpoIID (5), spatial confinement of the DMP complex by SpoIVA tracks would facilitate these functional interactions. In the absence of SpoIVA, DMP components may randomly disperse along the leading edge of the engulfing membrane, leading to inefficient PG hydrolysis and engulfment defects. Additionally, SpoIVA tracks may restrict broad diffusion of the DMP complex outside the engulfing membrane.

We a propose a model in which SpoIVA polymers provides a dual-function scaffold: 1) they promote forward movement of the engulfing membrane, and 2) the tracks they form organize DMP complex activity by corralling it to discrete sites at the leading edge of the engulfing membrane (Fig. 6). Testing the latter aspect of this model would require high-resolution, single-molecule microscopy approaches to analyse the diffusion of DMP complexes and this remains an important challenge for future work. Interestingly, examination of published TEM images of sporulating *Clostridioides difficile* cells revealed the presence of indented forespores with protrusions (41) similar to those reported in this study. Thus, SpoIVA may also play a similar role in engulfment in other spore-forming bacteria.

Overall, our data reveal new insight into the complexity of engulfment during spore development and pave the way for future studies aimed at deciphering the exact role of SpoIVA during engulfment in other spore-formers.

## ACKNOWLEDGEMENTS

We thank members of the Rodrigues laboratory, past and present, for their support and encouragement. We thank Danae Angeles Morales for construction of pDMA006 and Cerith Harries, and their media and reagent preparation room staff, for continued support of our work. We acknowledge the Midlands Regional Cryo-EM Facility, hosted at the Warwick Advanced Bioimaging Research Technology Platform, for use of the JEOL 2100Plus, supported by MRC award reference MC_PC_1713. This work was partly supported by grant BB/X008533/1 from the Biotechnology and Biological Sciences Research Council (https://www.ukri.org/councils/bbsrc) to C.D.A.R and by the Midlands Integrative Biosciences Training Partnership (https://warwick.ac.uk/fac/cross_fac/mibtp/) scholarship to K. C (grant number BB/T00746X/1) and B. F (grant number BB/T00746X/1). IBS acknowledges integration into the Interdisciplinary Research Institute of Grenoble (IRIG, CEA).

## MATERIALS AND METHODS

### General methods

All *Bacillus subtilis* strains used in this study are derived from the autotrophic strain 168. Deletion mutants were generated using isothermal assembly. Approximately 1500 bp regions upstream and downstream of the gene of interest, an antibiotic resistance cassette, and plasmid backbone were PCR-amplified. The four fragments were assembled using isothermal assembly to generate a knockout plasmid. The resulting construct was introduced into *Bacillus subtilis* using a one-step transformation method. Chromosomal integration of the knockout was confirmed by PCR using a primer within the original cloning region and a second primer located outside the integration site.

Sporulation was induced either by resuspension at 37°C following the Sterlini-Mandelstam protocol (42) or by nutrient depletion in a supplemented Difco sporulation medium (DSM) (ref). The DSM medium contained 8 g/L Bacto nutrient broth (Difco), 0.1% (wt/vol) KCl, 1 mM MgSO₄, 0.5 mM NaOH, 1 mM Ca(NO₃)₂, 0.01 mM MnCl₂, and 0.001 mM FeSO₄. Sporulation efficiency was assessed using a heat-resistance assay. Cultures grown in DSM for 30 hours at 37°C were subjected to heat treatment at 80°C for 20 minutes, and the number of heat-resistant colony-forming units (CFUs) was compared to that of the wild-type strain.

### Fluorescence microscopy

Sporulating cells were obtained at predetermined time points via resuspension method, 200 µl of culture was collected and pelleted via centrifugation. These were resuspended with 10 µl of 0.05 mM TMA-DPH [1-(4-trimethylammoniumphenyl)-6-phenyl-1,3,5-hexatriene p-toluenesulfonate]. Next, 2 µl of strained samples were spread on 2% (w/v) low melting point agarose pads prepared with resuspension media mounted on Gene Frames (Bio-Rad) and a coverslip placed on top of the Gene Frame.

Cells were imaged using Zeiss Axio Observer microscope fitted with a Plan-Apochromat 100×/1.4 Oil Ph3 objective lens and a Colibri 7 Type R[G/Y]CBV-UV fluorescent light source. Images were captured with an Axiocam 712 mono camera. The TMA-DPH membrane dye was excited using a Zeiss Axio 92HE filter with a 100 ms exposure time. CFP fluorescence was acquired using a Zeiss Axio 108HE filter with 300 ms exposure, and GFP fluorescence was captured using the same filter set with a 100 ms exposure.

### Image processing and statistics

Microscopy images were processed by adjusting the brightness and contrast using Image Fiji software (43). Sporulating cells that contained flat septa, septal membrane bulges, even and uneven migration of the engulfing membrane were manually quantified using the Cell Counter plugin in Fiji.

Engulfment completion was assessed based on the membrane stain intensity around the forespore. When stained with TMA-DPH, unfused membranes on cells that have not completed engulfment have higher florescence intensity than cells which have completed engulfment. The number of cells that completed or did not complete engulfment was manually counted using the Cell Counter plugin in Fiji (ref).

GFP-SpoIIP fluorescence intensity was analysed using the MicrobeJ plugin for Fiji software (43). First, background florescence was subtracted (Process > Subtract Background) to avoid false-positive signal. The “Bacteria” tab in MicrobeJ was configured to “Smoothed” to detect sporangia outlines from phase-contrast images. Parameters “Exclude on Edges,” “Shape descriptors,” and “Segmentation” were selected. To detect fluorescent GFP-SpoIIP foci for mean fluorescence intensity measurements, the “Maxima” tab on MicrobeJ was set to “Point”. Violin plots were created using MicrobeJ data into Superplots tool (44) available at https://huygens.science.uva.nl/SuperPlotsOfData/. To compare distributions between populations of sporulating wild-type and mutant cells, a nonparametric Kolmogorov-Smirnov test was carried out. Welch’s *t*-tests were conducted to compare the means between these populations.

### Transmission electron microscopy

10 ml of cell culture was collected at desired time points and pelleted at 13,000 rpm for 5 min, after which they were transferred into a primary fixative (4% glutaraldehyde in 0.1 M sodium cacodylate buffer) and incubated overnight at 4°C. Samples were then stained with 1% osmium tetroxide for 1 h, followed by washing. After stepwise dehydration 25%, 50%, 75% and 100% acetone, they were infiltrated with 50% resin for 1 h, followed by 100% resin for 24 hours. The Agar LV resin was cured at 60°C degrees overnight. After ultrathin sectioning on an RMC ultramicrotome sections were post-stained in 2% uranyl acetate for 2 min and 1.5% lead citrate for 1 min. Samples were imaged in a JEOL JEM2100Plus with Gatan OneView CMOS camera.

## REFERENCES

1. Errington J. 2003. Regulation of endospore formation in Bacillus subtilis. Nat Rev Microbiol 1:117–26.

2. Piggot PJ, Hilbert DW. 2004. Sporulation of Bacillus subtilis. Curr Opin Microbiol 7:579–86.

3. Ojkic N, Lopez-Garrido J, Pogliano K, Endres RG. 2016. Cell-wall remodeling drives engulfment during Bacillus subtilis sporulation. Elife 5.

4. Popham DL, Bernhards CB. 2015. Spore Peptidoglycan. Microbiology Spectrum 3:1–20.

5. Morlot C, Uehara T, Marquis KA, Bernhardt TG, Rudner DZ. 2010. A highly coordinated cell wall degradation machine governs spore morphogenesis in Bacillus subtilis. Genes Dev 24:411–22.

6. McPherson DC, Driks A, Popham DL. 2001. Two class A high-molecular-weight penicillin-binding proteins of Bacillus subtilis play redundant roles in sporulation. J Bacteriol 183:6046–53.

7. Driks A, Eichenberger P. 2016. The Spore Coat. Microbiology Spectrum 4:1–22.

8. McKenney PT, Driks A, Eichenberger P. 2013. The Bacillus subtilis endospore: assembly and functions of the multilayered coat. Nat Rev Microbiol 11:33–44.

9. Catalano FA, Meador-Parton J, Popham DL, Driks A. 2001. Amino acids in the Bacillus subtilis morphogenetic protein SpoIVA with roles in spore coat and cortex formation. J Bacteriol 183:1645–54.

10. Castaing JP, Nagy A, Anantharaman V, Aravind L, Ramamurthi KS. 2013. ATP hydrolysis by a domain related to translation factor GTPases drives polymerization of a static bacterial morphogenetic protein. Proceedings of the National Academy of Sciences of the United States of America 110:E151–E160.

11. McKenney PT, Eichenberger P. 2012. Dynamics of spore coat morphogenesis in Bacillus subtilis. Mol Microbiol 83:245–60.

12. Eichenberger P, Fujita M, Jensen ST, Conlon EM, Rudner DZ, Wang ST, Ferguson C, Haga K, Sato T, Liu JS, Losick R. 2004. The program of gene transcription for a single differentiating cell type during sporulation in Bacillus subtilis. PLoS Biol 2:e328.

13. Broder DH, Pogliano K. 2006. Forespore engulfment mediated by a ratchet-like mechanism. Cell 126:917–28.

14. Aung S, Shum J, Abanes-De Mello A, Broder DH, Fredlund-Gutierrez J, Chiba S, Pogliano K. 2007. Dual localization pathways for the engulfment proteins during Bacillus subtilis sporulation. Mol Microbiol 65:1534–46.

15. Meyer P, Gutierrez J, Pogliano K, Dworkin J. 2010. Cell wall synthesis is necessary for membrane dynamics during sporulation of Bacillus subtilis. Molecular Microbiology 76:956–970.

16. Chan H, Taib N, Gilmore MC, Mohamed AMT, Hanna K, Luhur J, Nguyen H, Hafiz E, Cava F, Gribaldo S, Rudner D, Rodrigues CDA. 2022. Genetic Screens Identify Additional Genes Implicated in Envelope Remodeling during the Engulfment Stage of Bacillus subtilis Sporulation. mBio 13:e0173222.

17. Broder DH, Pogliano K. 2006. Forespore Engulfment Mediated by a Ratchet-Like Mechanism. Cell 126:917–928.

18. Ramamurthi KS, Clapham KR, Losick R. 2006. Peptide anchoring spore coat assembly to the outer forespore membrane in Bacillus subtilis. Molecular Microbiology 62:1547–1557.

19. Ramamurthi KS, Lecuyer S, Stone HA, Losick R. 2009. Geometric cue for protein localization in a bacterium. Science 323:1354–7.

20. Ramamurthi KS, Losick R. 2008. ATP-Driven Self-Assembly of a Morphogenetic Protein in Bacillus subtilis. Molecular Cell 31:406–414.

21. Delerue T, Updegrove TB, Chareyre S, Anantharaman V, Gilmore MC, Jenkins LM, Popham DL, Cava F, Aravind L, Ramamurthi KS. 2024. Bacterial spore surface nanoenvironment requires a AAA+ ATPase to promote MurG function. Proc Natl Acad Sci U S A 121:e2414737121.

22. Ebmeier SE, Tan IS, Clapham KR, Ramamurthi KS. 2012. Small proteins link coat and cortex assembly during sporulation in Bacillus subtilis. Molecular Microbiology 84:682–696.

23. Ozin AJ, Henriques AO, Yi H, Moran CP, Jr. 2000. Morphogenetic proteins SpoVID and SafA form a complex during assembly of the Bacillus subtilis spore coat. J Bacteriol 182:1828–33.

24. Costa T, Isidro AL, Moran CP, Henriques AO. 2006. Interaction between coat morphogenetic proteins SafA and SpoVID. Journal of Bacteriology 188:7731–7741.

25. de Francesco M, Jacobs JZ, Nunes F, Serrano M, McKenney PT, Chua M-H, Henriques AO, Eichenberger P. 2012. Physical Interaction between Coat Morphogenetic Proteins SpoVID and CotE Is Necessary for Spore Encasement in Bacillus subtilis. Journal of Bacteriology 194:4941–4950.

26. Nunes F, Fernandes C, Freitas C, Marini E, Serrano M, Moran CP, Eichenberger P, Henriques AO. 2018. SpoVID functions as a non-competitive hub that connects the modules for assembly of the inner and outer spore coat layers in Bacillus subtilis. Molecular Microbiology 110:576–595.

27. Ozin AJ, Samford CS, Henriques AO, Moran CP. 2001. SpoVID guides SafA to the spore coat in Bacillus subtilis. Journal of Bacteriology 183:3041–3049.

28. Fimlaid KA, Jensen O, Donnelly ML, Siegrist MS, Shen A. 2015. Regulation of Clostridium difficile Spore Formation by the SpoIIQ and SpoIIIA Proteins. PLoS Genet 11:e1005562.

29. Luhur J, Chan H, Kachappilly B, Mohamed A, Morlot C, Awad M, Lyras D, Taib N, Gribaldo S, Rudner DZ, Rodrigues CDA. 2020. A dynamic, ring-forming MucB / RseB-like protein influences spore shape in Bacillus subtilis. PLoS Genet 16:e1009246.

30. Perez AR, Abanes-De Mello A, Pogliano K. 2000. SpoIIB localizes to active sites of septal biogenesis and spatially regulates septal thinning during engulfment in bacillus subtilis. J Bacteriol 182:1096–108.

31. Abanes-De Mello A, Sun YL, Aung S, Pogliano K. 2002. A cytoskeleton-like role for the bacterial cell wall during engulfment of the Bacillus subtilis forespore. Genes Dev 16:3253–64.

32. Gutierrez J, Smith R, Pogliano K. 2010. SpoIID-mediated peptidoglycan degradation is required throughout engulfment during Bacillus subtilis sporulation. J Bacteriol 192:3174–86.

33. Chastanet A, Losick R. 2007. Engulfment during sporulation in Bacillus subtilis is governed by a multi-protein complex containing tandemly acting autolysins. Mol Microbiol 64:139–52.

34. Mohamed AMT, Chan H, Luhur J, Bauda E, Gallet B, Morlot C, Cole L, Awad M, Crawford S, Lyras D, Rudner DZ, Rodrigues CDA. 2021. Chromosome Segregation and Peptidoglycan Remodeling Are Coordinated at a Highly Stabilized Septal Pore to Maintain Bacterial Spore Development. Dev Cell 56:36–51 e5.

35. Doan T, Morlot C, Meisner J, Serrano M, Henriques AO, Moran CP, Jr., Rudner DZ. 2009. Novel secretion apparatus maintains spore integrity and developmental gene expression in Bacillus subtilis. PLoS Genet 5:e1000566.

36. Rodrigues CD, Marquis KA, Meisner J, Rudner DZ. 2013. Peptidoglycan hydrolysis is required for assembly and activity of the transenvelope secretion complex during sporulation in Bacillus subtilis. Mol Microbiol 89:1039–52.

37. Rodrigues CD, Ramirez-Guadiana FH, Meeske AJ, Wang X, Rudner DZ. 2016. GerM is required to assemble the basal platform of the SpoIIIA-SpoIIQ transenvelope complex during sporulation in Bacillus subtilis. Mol Microbiol 102:260–273.

38. Mylvaganam S, Freeman SA, Grinstein S. 2021. The cytoskeleton in phagocytosis and macropinocytosis. Curr Biol 31:R619–R632.

39. Jaumouille V, Waterman CM. 2020. Physical Constraints and Forces Involved in Phagocytosis. Front Immunol 11:1097.

40. Freeman SA, Vega A, Riedl M, Collins RF, Ostrowski PP, Woods EC, Bertozzi CR, Tammi MI, Lidke DS, Johnson P, Mayor S, Jaqaman K, Grinstein S. 2018. Transmembrane Pickets Connect Cyto- and Pericellular Skeletons Forming Barriers to Receptor Engagement. Cell 172:305–317 e10.

41. Touchette MH, Benito de la Puebla H, Alves Feliciano C, Tanenbaum B, Schenone M, Carr SA, Shen A. 2021. Identification of a Novel Regulator of Clostridioides difficile Cortex Formation. mSphere 6:e0021121.

42. Sterlini JM, Mandelstam J. 1969. Commitment to sporulation in Bacillus subtilis and its relationship to development of actinomycin resistance. The Biochemical journal 113:29–37.

43. Schindelin J, Arganda-Carreras I, Frise E, Kaynig V, Longair M, Pietzsch T, Preibisch S, Rueden C, Saalfeld S, Schmid B, Tinevez JY, White DJ, Hartenstein V, Eliceiri K, Tomancak P, Cardona A. 2012. Fiji: an open-source platform for biological-image analysis. Nat Methods 9:676–82.

44. Goedhart J. 2021. SuperPlotsOfData-a web app for the transparent display and quantitative comparison of continuous data from different conditions. Mol Biol Cell 32:470–474.

45. E. Bauda, B. Fekade, L. Bellard, B. Gallet, S. Degroux, E. Neumann, C. Masc, K. Coleman, A. Leroy, G. Effantin, D. Fenel, J. Moravcova, J. Novacek, C. Moriscot, G. Schoehn, C. D.A. Rodrigues, and C. Morlot. 2026. Assembly of the Essential SpoIVA Coat Protein at the Surface of Developing Bacterial Spores. BioRxiv.

