## Supporting Figures for "SpoIVA contributes to efficient engulfment through a cytoskeletal-like mechanism during *Bacillus subtilis* sporulation"

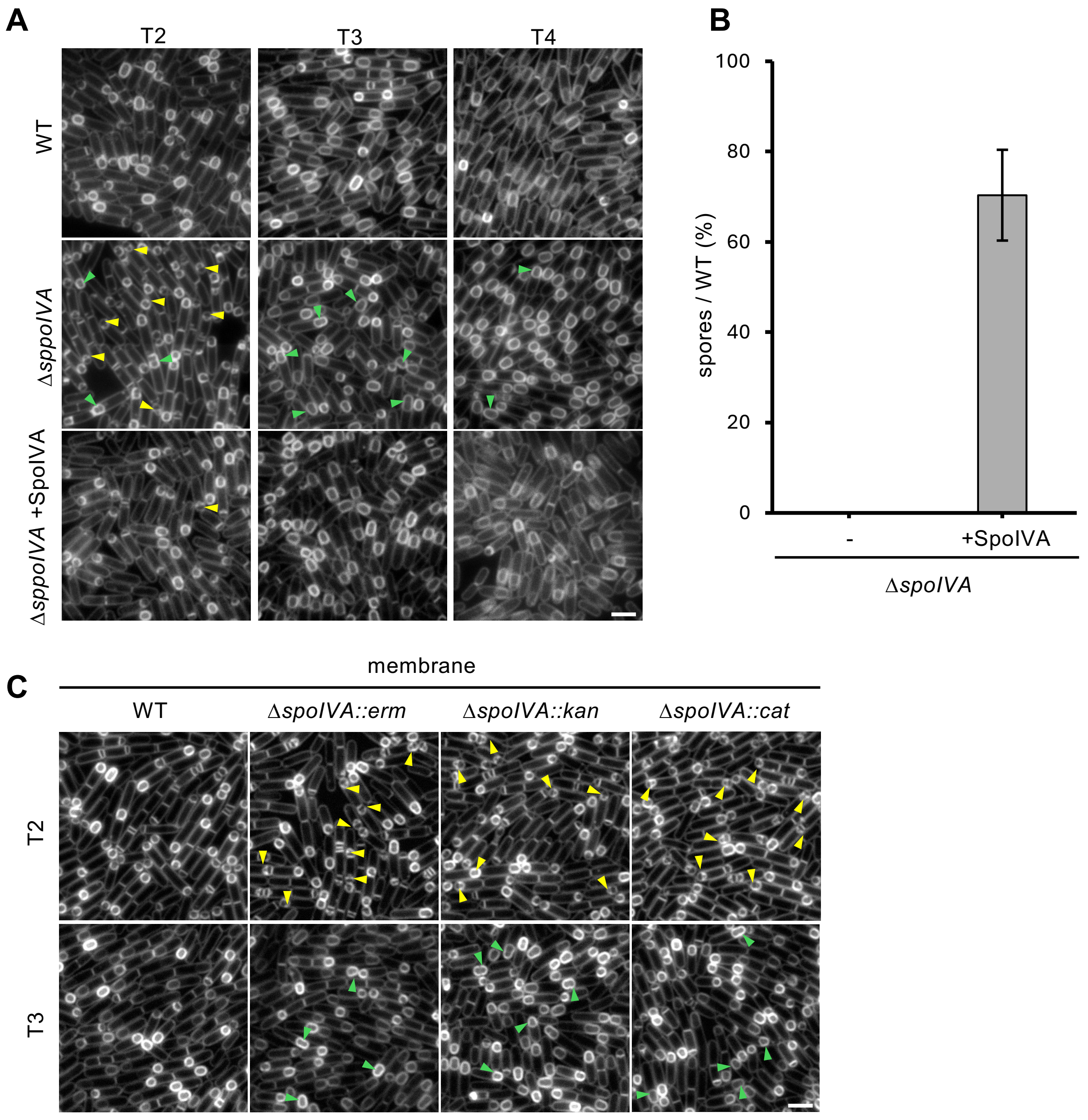


**Figure S1. Complementation of SpoIVA from an ectopic locus. (A)** Representative images of sporulating cells at 2 hours (T2), 3 hours (T3) and 4 hours (T4) in the wild-type (WT) and Δ*spoIVA* and SpoIVA complementation strains. Forespore cytoplasm were visualised using a forespore reporter ($P_{spoIIQ}$-*cfp*) that appears cyan in merged images. Cell membranes false-coloured red in merged images. Septal membrane bulges are highlighted with yellow triangles and indented cells are highlighted with green triangles. **(B)** Δ*spoIVA* phenotype can be complemented by expression of SpoIVA at an ectopic locus. Shown is the mean percentage (mean percentage ± SD, n=3) of heat-resistant cells after sporulation by nutrient exhaustion in the presence and absence of SpoIVA as a percentage of WT. SpoIVA complementation restores sporulation efficiency to 70% ± 20%. Error bars represent SD from three biological replicates. **(C)** Representative images of sporulating cells at 2 hours (T2), 3 hours (T3) and 4 hours (T4) in Δ*spoIVA* strains show that spore defects are not marker-specific. Scale bar = 2 µm.

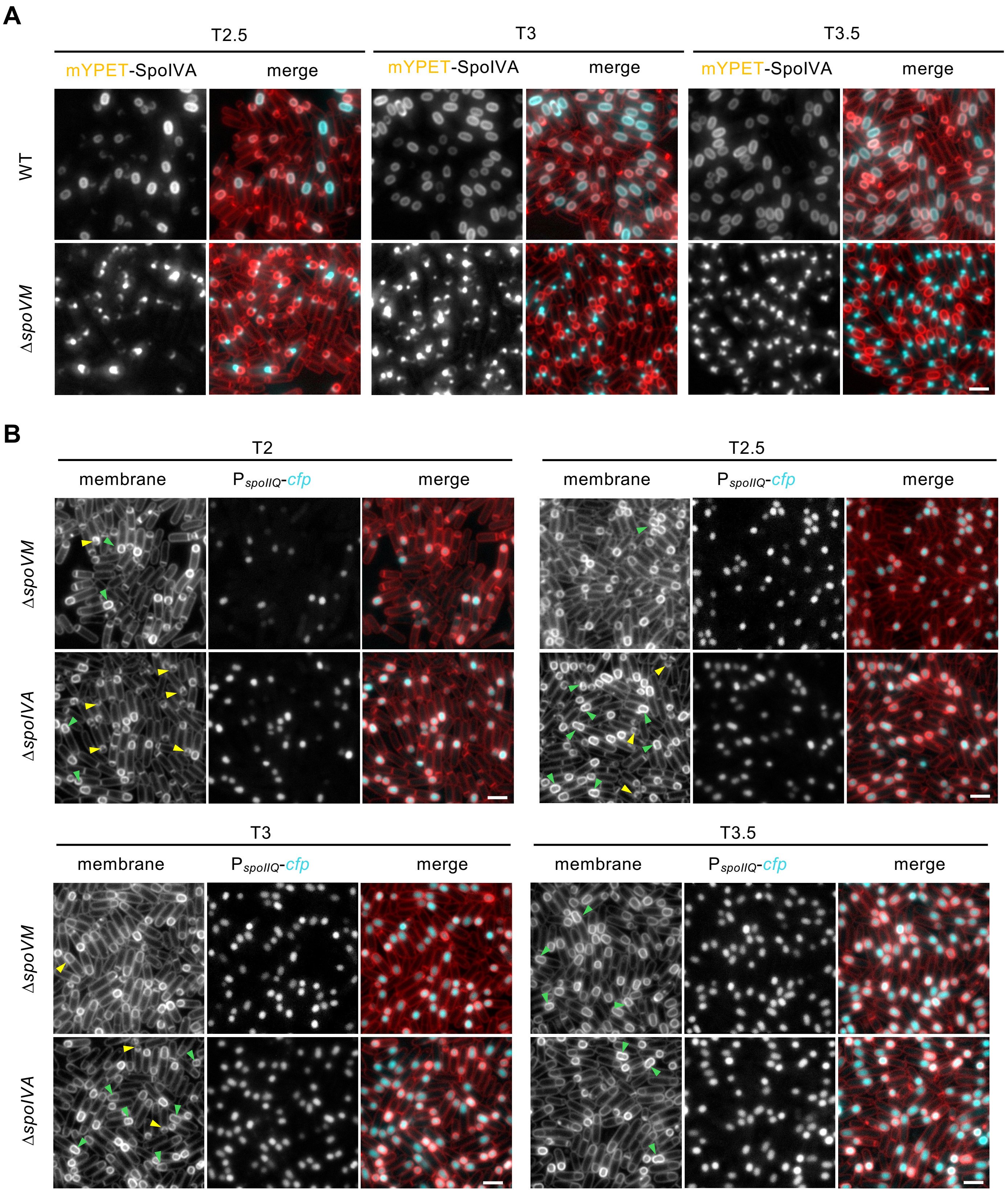


**Figure S2. Localization of SpoIVA in absence of SpoVM and engulfment defects in the Δ*spoVM* mutant. (A)** mYPET-SpoIVA localization in wild-type (Wt), Δ*spoVM* strain at 2.5 hours (T2.5) and 3 hours (T3.5) after the onset of sporulation. **(B)** Engulfment progression. In the Δ*spoVM* and Δ*spoIVA* strains at 2 hours (T2), 2.5 hours (T2.5) and 3 hours (T3.5) after the onset of sporulation. Forespore cytoplasm were visualised using a forespore reporter ($P_{spoIIQ}$-*cfp*) that appears cyan in merged images. Cell membranes are false-coloured red in merged images. Septal membrane bulges are highlighted with yellow triangles and indented cells are highlighted with green triangles. Scale bar = 2 μm.

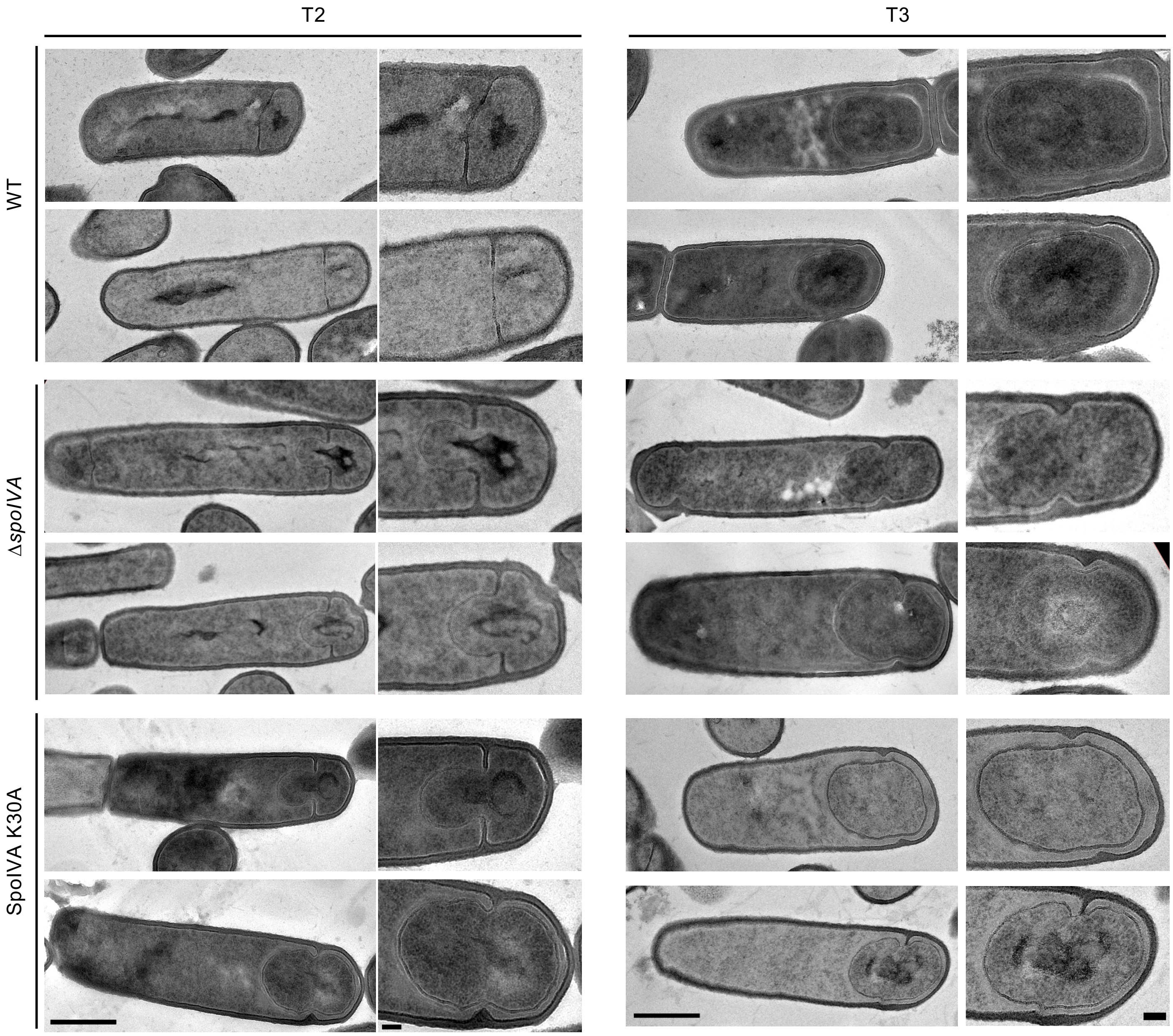


**Figure S3. Electron micrographs of in sporulating cells.** Representaitve images of sporulating cells in the wild-type (WT), Δ*spoIVA* and SpoIVA K30Aa mutants at 2 hours (T2) and 3 hours (T3) after the onset of sporulation. Δ*spoIVA* and SpoIVA K30A mutants show bulging phenotype at T2 and indented phenotype after engulfment completion at T3. Scale bar is 500 nm and 100 nm in standard and zoomed-in images, respectively.

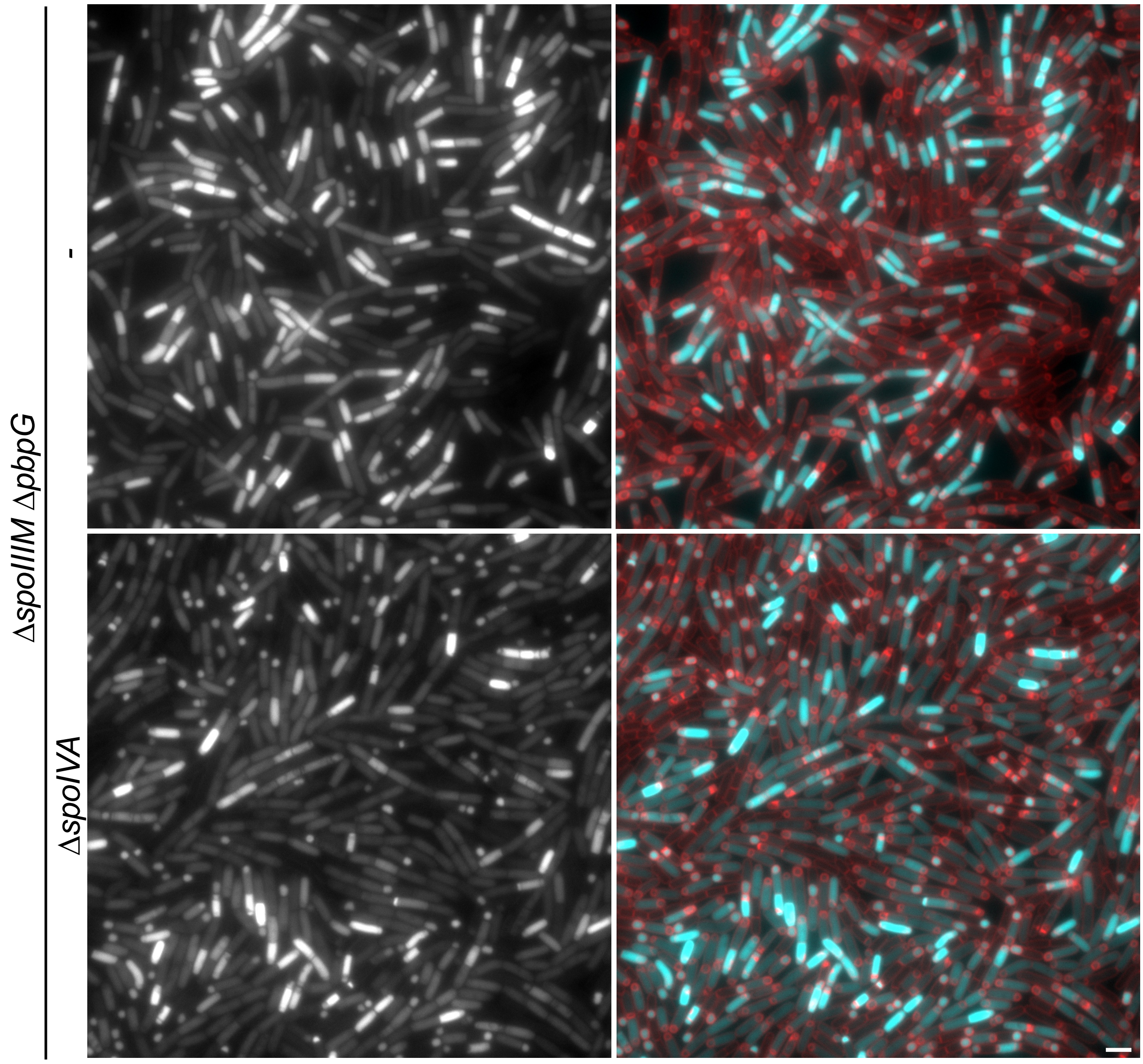


**Figure S4. The Δ*spoIVA* mutant increases compartmentalisation defect of SpoIIIM and PbpG mutants.** Large panel representative images of miscompartmentalisation in wild-type (WT), Δ*spoIIIM* Δ*pbpG* and Δ*spoIIIM* Δ*pbpG* Δ*spoIVA* strains at 3 hours (T3) after the onset of sporulation. Forespore cytoplasm were visualised using a forespore reporter ($P_{spoIIQ}$-cfp) that appears cyan in merged images. Cell membranes are false-coloured red in merged images. Scale bar = 2 µm.

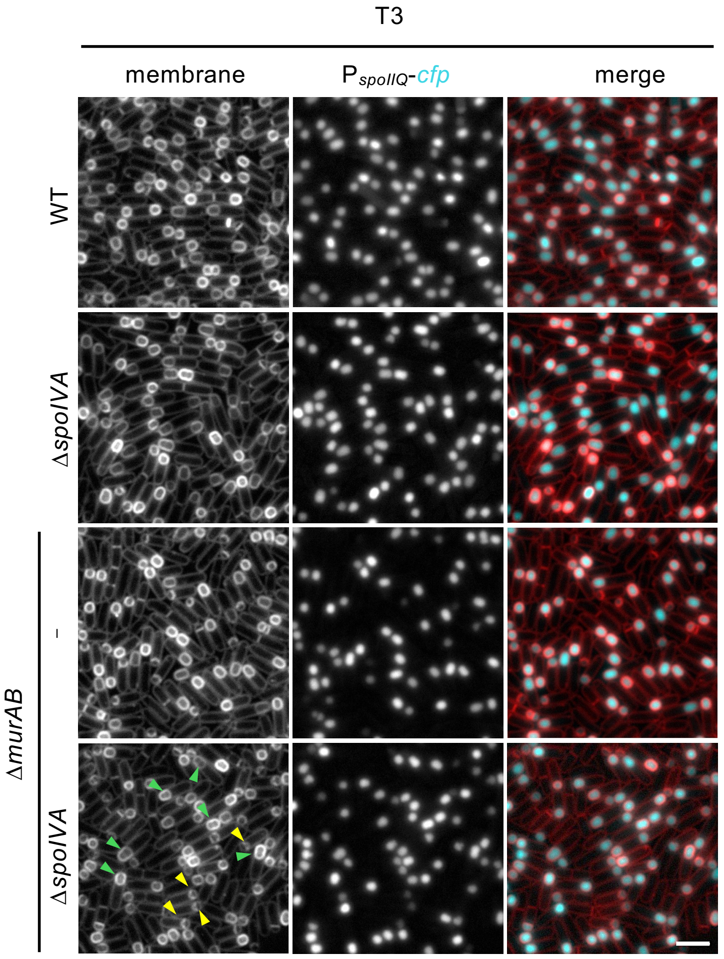


**Figure S5. Engulfment defects in the Δ*spoIVA*, Δ*murAB* and Δ *spoIVA* Δ*murAB* mutant.** Engulfment progression in the wild-type (WT), Δ*spoIVA*, Δ*murAB and* Δ*spoIVA* Δ*murAB* strains at 3 hours (T2) after the onset of sporulation. Forespore cytoplasm were visualised using a forespore reporter ($P_{spoIIQ}$-cfp) that appears cyan in merged images. Cell membranes are false-coloured red in merged images. Septal membrane bulges are highlighted with yellow triangles and indented forespores are highlighted with green triangles. Scale bar = 2 µm.

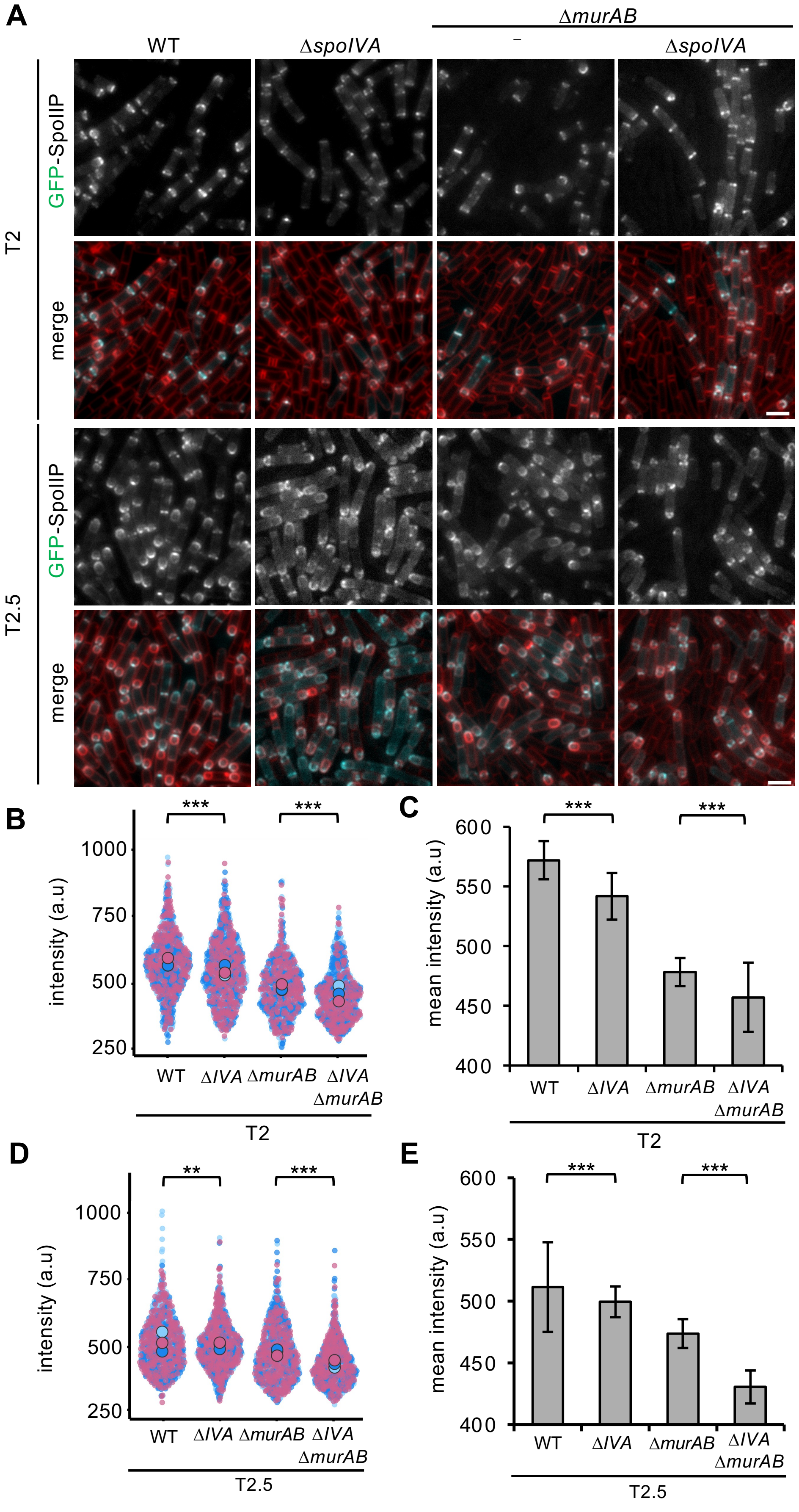


**Figure S6. GFP-SpoIIP localization in the absence of *spoIVA* and *murAB*. (A)** GFP-SpoIIP localisation at T2 and T2.5. Cell membranes and GFP are false-coloured red and cyan in merged images. Scale bar = 2 µm. **(B)** Distribution of GFP-SpoIIP intensity. Superplots contain data from 3 biological replicates with a total of 795, 735, 645 and 694 cells for WT, Δ*spoIVA*, Δ*murAB* and Δ*spoIVA* Δ*murAB,* respectively, at T2. **(C)** mean GFP-SpoIIP intensity (a.u.) at 2 hours (T2) after onset of sporulation. **(D)** Distribution of GFP-SpoIIP intensity. Superplots contain data from 3 biological replicates with a total number of 713, 822, 742 and 743 cells for WT, Δ*spoIVA*, Δ*murAB* and Δ*spoIVA* Δ*murAB,* respectively, at T2.5. **(E)** Mean GFP-SpoIIP intensity (a.u.) at 2.5 hours (T2.5) after onset of sporulation in the wild-type, Δ*spoIVA*, Δ*murAB* and Δ*spoIVA* Δ*murAB* strains. For panel B & D, ** $p < 0.01$ and *** $p < 0.001$ using Kolmogorov-Smirnov test to compare total population distributions. For panel C & E *** $p < 0.001$ by Welch's t test performed on the mean intensity, replicates (n=3) at T2 and T2.5 for the WT versus Δ*spoIVA,* Δ*murAB* and Δ*spoIVA* Δ*murAB* mutant.

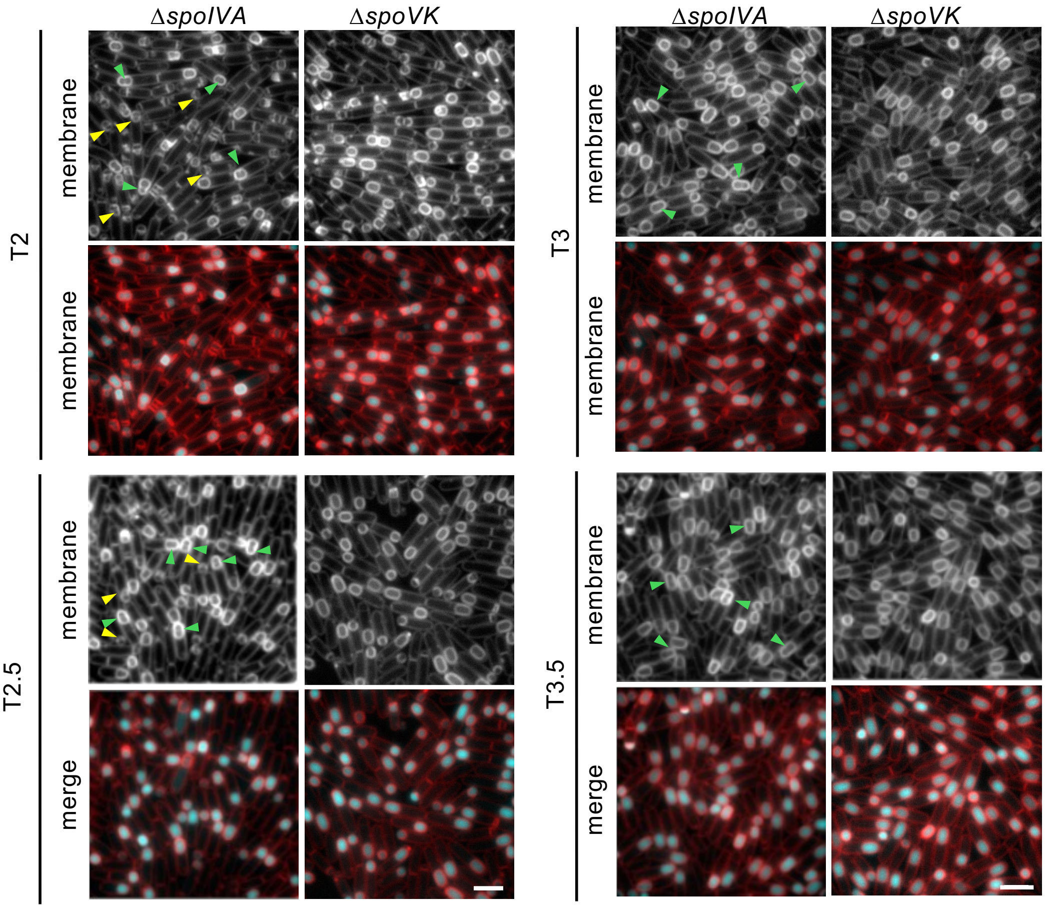


**Figure S7. Engulfment defects in Δ*spoIVA* mutants are not caused by SpoVK mislocalisation** Representative images of sporulating cells at 2 hours (T2), 2.5 hours (T2.5), 3 hours (T3) and 3.5 hours (T3.5) in the wild-type (WT) and Δ*spoIVA* and Δ*spoVK* mutants. Forespore cytoplasm were visualised using a forespore reporter ($P_{spoIIQ}$-cfp) that appears cyan in merged images. Cell membranes are false-coloured red in merged images. Septal membrane bulges are highlighted with yellow triangles and indented forespores are highlighted with green triangles. Scale bar = 2 µm.

Fekade *et al.*, Figure S8

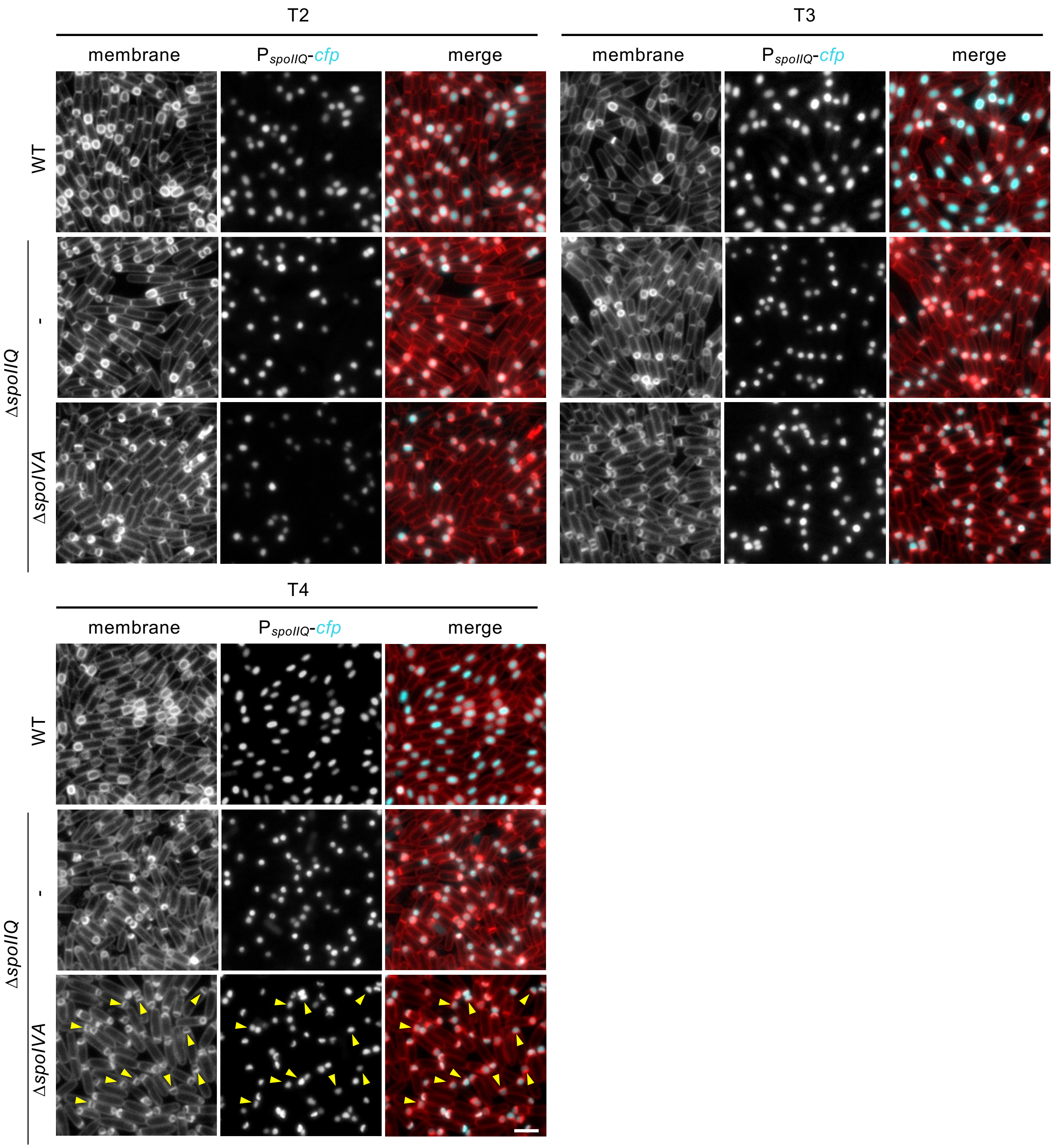


**Figure S8. Engulfment is severely impaired in ΔspoIIQ ΔspoIVA double mutants. (A)** Representative images of sporulating cells at 2 hours (T2), 3 hours (T3) and 4 hours (4) in the wild-type (WT) and Δ*spoIIQ* and Δ*spoIIQ* Δ*spoIVA* mutants. Forespore cytoplasm were visualised using a forespore reporter ($P_{spoIIQ}$-cfp) that appears cyan in merged images. Cell membranes are false-coloured red in merged images. Scale bar = 2 µm.

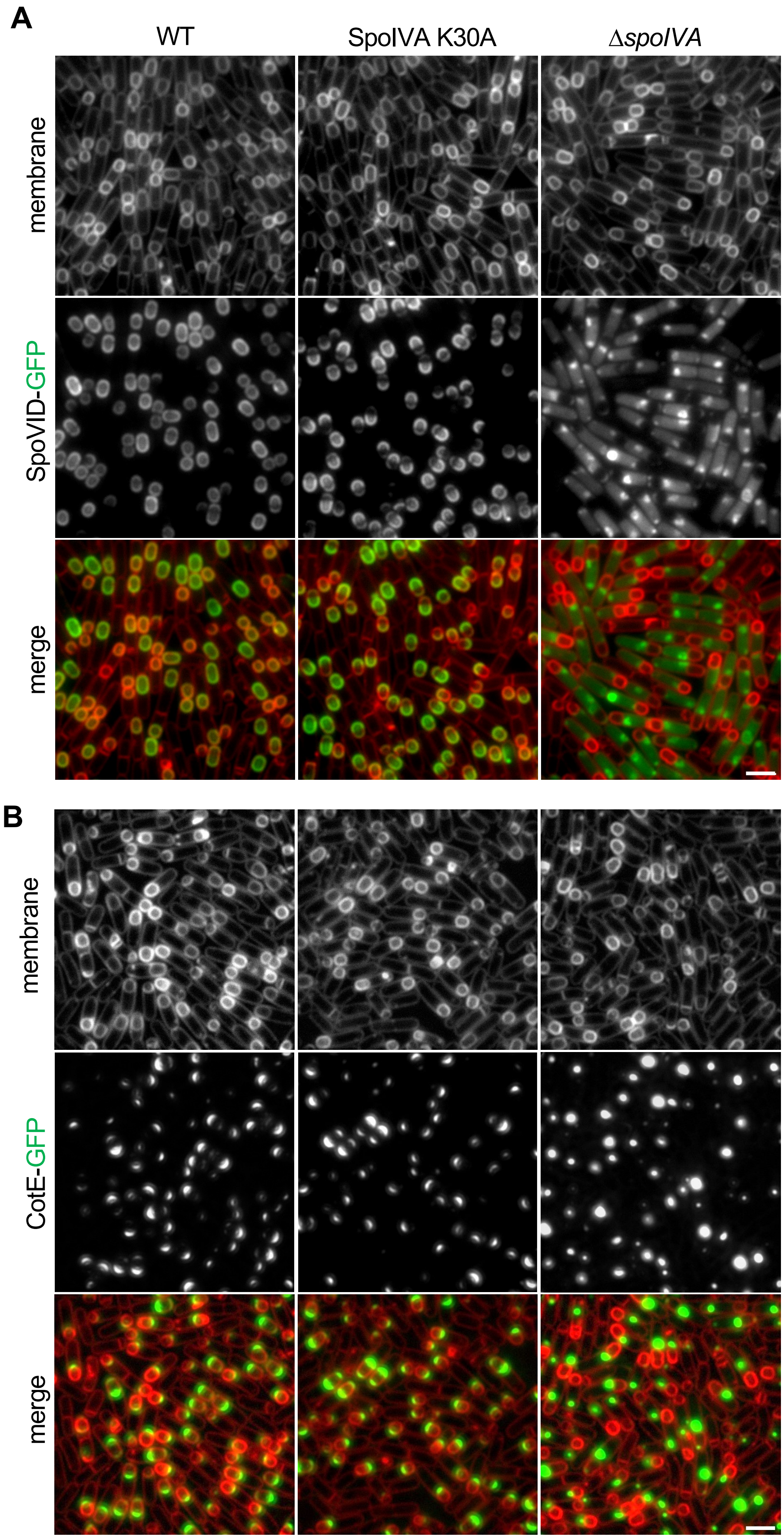


**Figure S9. SpoVID-GFP and CotE-GFP remains localized in the SpoIVA K30A mutant.** Representative images of **(A)** SpoVID-GFP and (**B)** CotE-GFP localization in the Δ*spoIVA* and SpoIVA K30A at T3. Cell membranes were visualized with TMA-DPH fluorescent membrane dye and are false-coloured red in merged images. Scale bar = 2 μm.

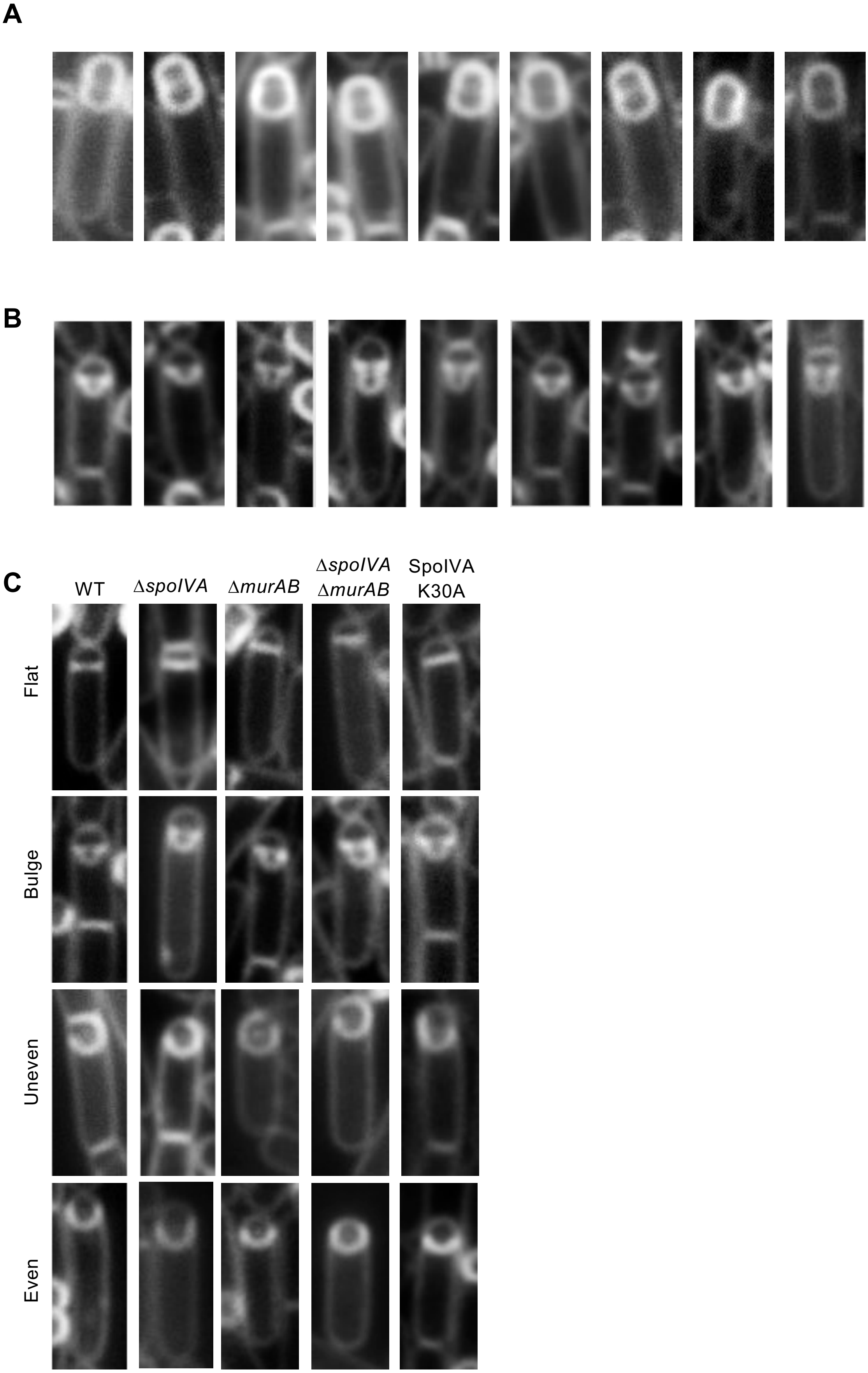


**Figure S10. Representative images of engulfment defect in WT and various mutants. (A)** Representative images of indented phenotype in Δ*spoIVA* mutants at at 2 hours (T2) after onset of sporulation. **(B)** Representative images of bulge phenotype in Δ*spoIVA* mutants at at 2 hours (T2) after onset of sporulation. **(C)** Representative membrane fluorescence images of flat, bulge, uneven and even septa in sporulating WT, Δ*spoIVA*, Δ*murAB*, Δ*spoIVA* Δ*murAB*, and SpoIVA K30A cells. Cell membranes were visualised using the fluorescent membrane dye.
