## Supporting Information for "SpoIVA contributes to efficient engulfment through a cytoskeletal-like mechanism during *Bacillus subtilis* sporulation"

**Table S1:** *Bacillus subtilis* strains used in this study.

| Strain | Genotype | Source |
|---|---|---|
| 168 | *auxotrophic wild-type strain* | Zeigler *et al.*, 2008 |
| bAT068 | *lacA::PspoIIQ-cfp (erm)* | This work |
| bAT087 | *amyE::PspoIIQ-cfp (cat)* | Mohamed *et al.*, 2021 |
| bAT091 | *amyE::PspoIIQ-cfp (cat), pbpG::kan, spoIIIM::erm* | Mohamed *et al.*, 2021 |
| bAT204 | *spoIIP::tet Ω PspoIIP-gfp–spoIIP (erm)* | This work |
| bAT478 | *amyE::PspoIIQ-cfp (cat, spoIIQ::erm* | Mohamed *et al.*, 2021 |
| bBD017 | *ycgO::spoIIE D584A (phleo), spoIIIE::lox72, amyE::PspoIIQ-cfp (cat),* | Dehghani *et al.*, 2024 |
| bBF334 | *lacA::PspoIIQ-cfp (erm), spoIVA::spec* | This work |
| bBF335 | *lacA::PspoIIQ-cfp (erm), spoVK::spec* | This work |
| bBF424 | *ycgO::spoIIIE D584A (phleo), spoIIIE::lox72, amyE::PspoIIQ-cfp (cat), spoIVA::erm* | This work |
| bBF426 | *spoIVA::spec, ycgO::PspoIVA-mYPET-spoIVA (cat), spoVM::erm* | This work |
| bBF496 | *cotEΩcotE-gfp (spec), spoIVA::erm* | This work |
| bBF497 | *cotEΩcotE-gfp (spec), spoIVA::erm, ycgo::PspoIVA-spoIVA (K30A)* | This work |
| bDMA099 | *spoIVA::lox72* | This work |
| bDMA125 | *spoIVA::lox72, yhdG::PspoIVA-spoIVA (tet)* | This work |
| bHC225 | *spoVIDΩspoVID-gfp (spec), spoIVA::cat* | This work |
| bHC332 | *spoIVA::erm* | This work |
| bHC405 | *spoIVA::spec, ycgO::PspoIVA-spoIVA (K30A) (cat)* | This work |
| bHC483 | *spoIVA::kan* | This work |
| bHC504 | *spoVIDΩspoVID-gfp (spec), spoIVA::kan, ycgO::PspoIVA-spoIVA (K30A) (cat)* | This work |
| bJL011 | *cotEΩcotE-gfp (spec)* | This work |
| bJL012 | *spoVIDΩspoVID-gfp (spec)* | Luhur *et al.*, 2020 |
| bJL043 | *spoIVA::cat* | Luhur *et al.*, 2020 |
| bJL069 | *lacA::PspoIIQ-cfp(erm), spoIVA::cat* | This work |
| bJL129 | *ycgO::PspoIVA-mYPET-spoIVA (cat)* | This work |
| bJL173 | *spoIVA::spec* | This work |
| bSG009 | *lacA::PspoIIQ-cfp (erm), spoIVA::spec, ycgO::PspoIV-spoIVA (K30A) (cat)* | This work |
| bSG010 | *spoIIP::tet Ω PspoIIP-gfp–spoIIP (erm), spoIVA::spec* | This work |
| bSG014 | *amyE::PspoIIQ-cfp (cat), pbpG::kan, spoIIIM::erm, spoIVA::spec* | This work |
| bSG015 | *lacA::PspoIIQ-cfp (erm), murAB::kan* | This work |
| bSG016 | *lacA::PspoIIQ-cfp (erm), spoIVA::cat, murAB::kan* | This work |
| bSG021 | *spoIIP::tet Ω PspoIIP-gfp–spoIIP (erm), murAB::kan* | This work |
| bSG022 | *spoIIP::tet Ω PspoIIP-gfp–spoIIP (erm), spoIVA::spec, murAB::kan* | This work |
| bSG031 | *spoVM::erm, yvbJ::PspoIIQ-cfp (spec)* | This work |
| bSG032 | *spoVM::erm, yvbJ::PspoIIQ-cfp (spec), spoIVA::kan* | This work |
| bSG033 | *spoVM::erm, yvbJ::PspoIIQ-cfp (spec), spoIVA::kan, ycgO::PspoIVA-spoIVA (K30A) (cat)* | This work |
| bSG034 | *spoIIP::tet Ω PspoIIP-gfp–spoIIP (erm), spoIVA::spec, ycgO::PspoIVA-spoIVA (K30A) (cat)* | This work |
| bSG035 | *spoIIP::tet Ω PspoIIP-gfp–spoIIP (erm), spoIVA::spec, murAB::kan ycgO::PspoIVA-spoIVA (K30A) (cat)* | This work |
| bSG036 | *amyE::PspoIIQ-cfp (cat), spoIIQ::erm, spoIVA::spec* | This work |

**Table S2:** Plasmids used in this study.

| Plasmid | Description | Source |
|---|---|---|
| pBF021 | *spoVK::spec* | This work |
| pDMA006 | *yhdG::PspoIVA-spoIVA (tet)* | This work |

### Plasmid Construction

**pDMA006 [*yhdG::PspoIVA-spoIVA (tet*)]** was generated via a two-way ligation containing *Bam*HI-*XhoI* PCR product of *PspoIVA-spoIVA* (oligonucleotides oCR740 and oDMA061) and plasmid pBB281 (*yhdG::tet*) cut with *Bam*HI and *XhoI*. pBB281 is an ectopic integration vector for double cross over integration a the non-essential *yhdG* locus (David Rudner laboratory stock).

**pBF021 [*spoVK::spec*]** was generated via a four-way ligation containing PCR products containing a 1500 bp DNA fragment upstream of *spoVK* (oligonucleotides oBF038 and oBF039), a spectinomycin resistance cassette (oligonucleotides oCR624 and oCR625), a 1500 bp fragment downstream of *spoVK* (oligonucleotides oBF040 and oBF041), and the vector backbone derived from pBB281 (oligonucleotides oBF069 and oBF070). The upstream fragment, downstream fragment, and vector backbone were each designed with 21 bp overlapping regions to enable efficient ligation.

**Table S3:** Oligonucleotide primers used in this study.

| Oligonucleotide | Sequence* |
|---|---|
| oCR740 | cgcGGATCCttacaggatgatggcgattaagcc |
| oDMA061 | ggcCTCGAGttggaaaaggtcgatattttcaagga |
| oBF038 | acatggtcacagtttgttgttgata |
| oBF039 | gtactgagcgagggagcagaagtccttcacctctctgttctc |
| oCR624 | ttctgctccctcgctcag |
| oCR625 | cagggagcactggtcaac |
| oBF040 | gtagttgaccagtgctccctgggaacctctcagtttttgaga |
| oBF041 | ctatttgtttatgaatgccggcaatg |
| oBF042 | ggtctgaggacaccgaacaaagc |
| oBF069 | ccggcattcataaacaaatagtctagagggaaaccgttgtgg |
| oBF070 | tcaacaacaaactgtgaccatgctgagcaataactagcataa |

*capital letters indicate restriction sites

### REFERENCES

A Mohamed, H Chan, J Luhur, E Bauda, B Gallet, C Morlot, L Cole, M Awad, S Crawford, D Lyras, DZ Rudner, CDA Rodrigues (2021) Chromosome segregation and peptidoglycan remodeling are coordinated at a highly stabilized septal pore to maintain bacterial spore development. *Developmental Cell* 11; 56 (1): 36-51.

B Dehghani, CDA Rodrigues (2024) SpoIIQ-dependent localization of SpoIIE contributes to septal stability and compartmentalization during the engulfment stage of Bacillus subtilis sporulation. *Journal of Bacteriology* 206:e00220-24.

J Luhur, H Chan, B Kachappilly, A Mohamed, C Morlot, M Awad, D Lyras, N Taib, S Gribaldo, DZ Rudner, CDA Rodrigues (2020) A dynamic, ring-forming MucB/RseB-like protein influences spore shape in *Bacillus subtilis*. *PLoS Genetics* 16(12): e1009246.

Zeigler DR, Prágai Z, Rodriguez S, Chevreux B, Muffler A, et al. (2008) The origins of 168, W23, and other *Bacillus subtilis* legacy strains. *J Bacteriol* 190: 6983–95. pmid:18723616
